# Autism-associated *Scn2a* haploinsufficiency disrupts *in vivo* dendritic signaling and impairs flexible decision-making

**DOI:** 10.1101/2025.04.08.647866

**Authors:** Hao Wu, Luqun Shen, Jonathan Indajang, Neil K. Savalia, Timothy G. Johnson, Jiayin Qu, Kevin J. Bender, Alex C. Kwan

## Abstract

*SCN2A* is a high-confidence risk gene for autism spectrum disorder. Loss-of-function mutations in *Scn2a* reduce dendritic excitability in neocortical pyramidal cells. However, the impact of *Scn2a* haploinsufficiency on dendritic signaling *in vivo*, particularly during behavior, is unknown. In this study, we used two-photon microscopy to image dendritic calcium transients in deep layer pyramidal cells in the mouse medial frontal cortex. *Scn2a^+/-^* mice had diminished coupling between apical and proximal dendritic compartments. Pyramidal tract neurons had abnormal event rates, while intratelencephalic neurons had compartment-specific alterations indicative of diminished dendritic integration. In a matching pennies task, *Scn2a^+/-^* mice were inflexible in the face of changing competitive pressure. Apical dendritic tuft in IT neurons typically encoded reward and strategy, but these task-specific representations were altered in *Scn2a^+/-^* mice. Collectively, the findings demonstrate that *Scn2a* haploinsufficiency weakens dendritic integration *in vivo* and disrupts the dendritic encoding of decision variables, potentially contributing to the cognitive rigidity in autism spectrum disorder.

## INTRODUCTION

Autism spectrum disorder (ASD) is a neurodevelopmental condition characterized by deficits in perception, social interaction, and cognition. At the genetic level, studies of rare coding variants have identified numerous high-confidence risk genes for ASD, many of which are involved in signaling and scaffolding in the postsynaptic dendritic compartments (1–3). Although the etiology of ASD remains poorly understood, growing evidence implicates deep layer glutamatergic neurons in the neocortex as key contributors (4). Alterations in excitatory transmission have been observed in many ASD models (5), and the synaptic dysfunctions may be closely tied to abnormalities in dendritic integration (6). Supporting this connection, structural alterations and calcium signaling perturbations to dendrites and dendritic spines have been reported in animal models for ASD (7–9) or autism related disorders such as Fragile X and Angelman syndrome (10, 11). However, uncovering the precise mechanisms by which the genetic perturbations lead to dendritic dysfunctions and ultimately behavioral deficits, remain an ongoing challenge.

Among the most strongly implicated high-confidence risk genes for ASD is *SCN2A* (2, 3). Loss-of-function mutations in *SCN2A* are associated with ASD and intellectual disability, whereas other alterations including gain-of-function mutations are more commonly associated with early onset epilepsy (12–14). *SCN2A* encodes Na_v_1.2, the type II alpha subunit of a voltage-gated sodium channel found in many cell types and brain regions. In the excitatory pyramidal cells in the neocortex, Na_v_1.2 is initially localized to the axon initiation segment during early development, but later becomes the dominant isoform expressed in the somatodendritic region in adulthood (15, 16). A function of Na_v_1.2 is to facilitate membrane excitability. Indeed, in mice with *Scn2a* haploinsufficiency, a disruption of Na_v_1.2 expression reduces dendrite excitability and impairs synaptic plasticity (15). Specifically, when the cell body is stimulated, calcium influx associated with the backpropagation of action potentials is diminished, reflecting as a decrease in the amplitude in the apical dendritic shaft, and inability to invade into the distal dendritic tuft (15). While these results highlight the importance of *Scn2a* for dendritic function, the studies were performed in acute brain slices. The extent to which these deficits may manifest *in vivo* and their behavioral relevance are open questions.

Recent advances in two-photon microscopy have provided powerful new approaches for visualizing calcium signals in the dendrites of cortical pyramidal neurons *in vivo*. The optimization of genetically encoded calcium indicators has yielded reporters with high signal-to-noise ratios, enabling imaging in subcellular compartments of neurons in live mice (17). This improvement has led to exciting discoveries about how synaptic inputs may be propagated and integrated in dendritic compartments. For example, studies have mapped the feature selectivity of inputs impinging at different dendritic locations (18–21), and revealed choice- and reward-related dendritic signals in the sensorimotor cortex during decision-making tasks (22–26). A particularly valuable innovation is the development of multi-plane imaging. This technique relies on rapid axial focusing via one of several possible methods such as electrically tunable lens, acousto-optic deflectors, and remote focusing. Multi-plane imaging enables near simultaneous recording of fluorescence transients at several depths, which is especially useful for quantifying the coupling between multiple compartments of a neuron including various dendritic branches and the soma *in vivo* (22, 27, 28).

The goal of this study is to determine the effects of *Scn2a* haploinsufficiency on dendritic calcium signaling *in vivo*. Performing multi-plane imaging using an electrically tunable lens, we visualized calcium transients in the distal apical tuft, apical trunk, and proximal trunk of pyramidal neurons in the mouse medial frontal cortex. We found reduced coupling of dendritic calcium signals in *Scn2a^+/-^* animals relative to controls. To assess how *Scn2a* haploinsufficiency affects behavior, we developed a competitive decision-making task for head-fixed mice that encourages adaptive strategy switching. *Scn2a^+/-^* mice were cognitively rigid, failing to adjust to a computer opponent that changes its strategy. During this task, apical dendritic tuft in the frontal cortex normally encoded reward- and strategy-related signals, but these representations were disrupted in *Scn2a^+/-^* mice. Overall, the results suggest the *in vivo* dendritic functions in the medial frontal cortex are impaired by *Scn2a* haploinsufficiency, including during flexible decision-making.

## RESULTS

### Multi-plane imaging of dendritic calcium transients in frontal cortical PT and IT neurons

To study *Scn2a* haploinsufficiency, we compared heterozygous *Scn2a^+/-^* mice to littermate *Scn2a^+/+^* controls. Our goal was to characterize dendritic calcium transients in pyramidal cells in the medial frontal cortex (**Fig. 1A**). There are two major subpopulations of pyramidal cells: pyramidal tract (PT) and intratelencephalic (IT) neurons. PT and IT neurons have distinct features; they have different morphological characteristics, physiological properties, and long-range projection targets (29–31). Moreover, PT and IT neurons may have differential contributions to neuropsychiatric conditions and their treatments (32–34). To target a sparse number of PT neurons, we injected a low titer of AAVrg-hSyn-Cre into the ipsilateral pons, and AAV-CAG-Flex-GCaMP6f in the dorsal medial frontal cortex including the anterior cingulate cortex (ACAd) and medial premotor cortex (MOs) (**Fig. 1B**). Fluorescence images of histological coronal sections confirmed a sparse labeling of neurons restricted to the deep layers of the medial frontal cortex, consistent with the expected laminar distribution of PT neurons (**Fig. 1B**).

**Fig. 1.**
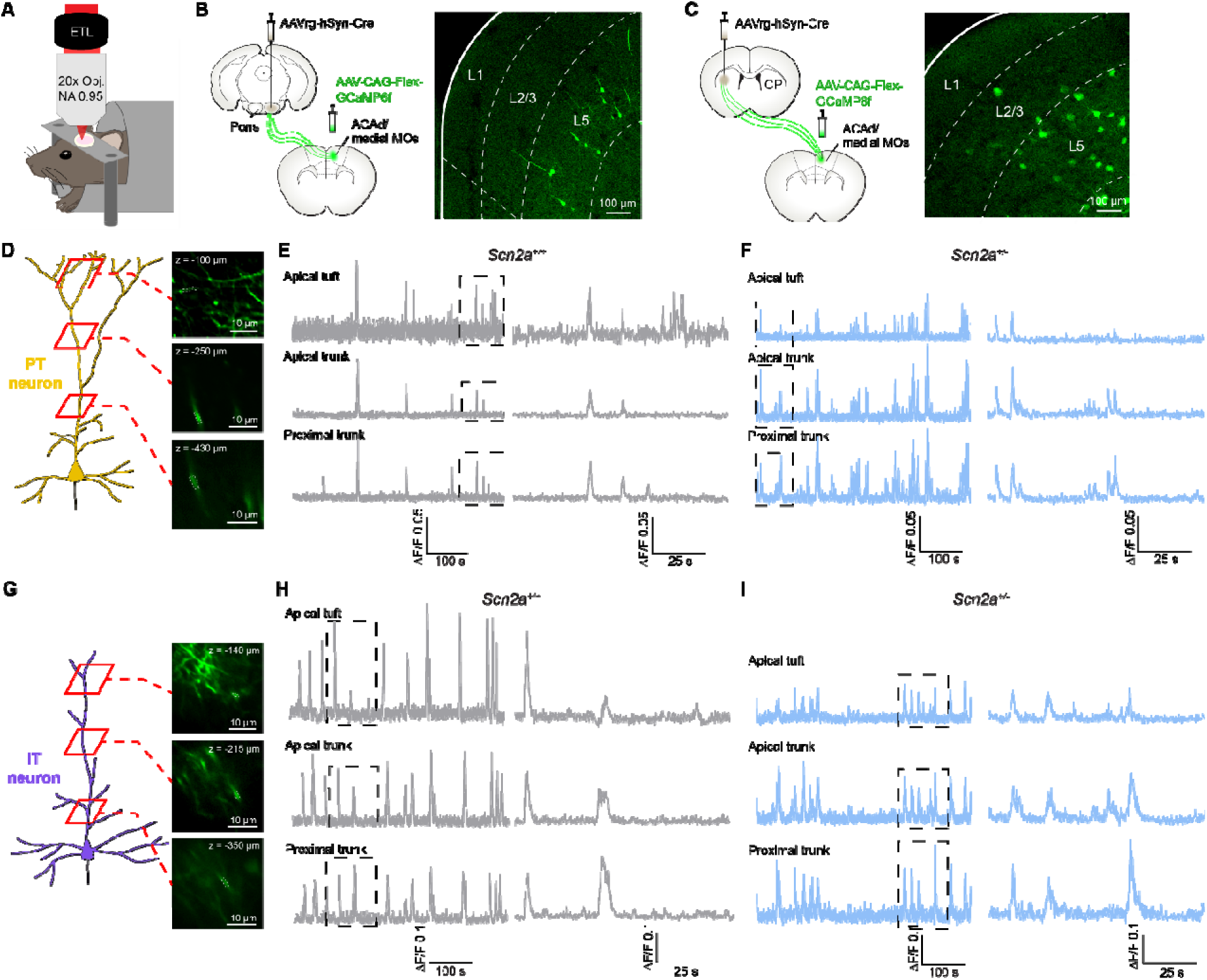
Imaging calcium transients in the dendritic compartments of single frontal cortical PT or IT neuron. **(A)** The imaging setup involving two-photon imaging of an awake, head-fixed mouse. ETL, electrically tunable lens. **(B)** Left, viral strategy to express GCaMP6f in frontal cortical PT neurons. Right, fixed coronal section showing GCaMP6f-expressing cells in the medial frontal cortex. White overlay, wireframe from the Allen Mouse Brain Atlas. **(C)** Similar to (B) for frontal cortical IT neurons. **(D)** Example two-photon images at different depths corresponding to apical tuft, apical trunk and proximal trunk of a frontal cortical PT neuron in a *Scn2a^+/+^* control mouse. Dash lines indicate the region of interest. **(E)** ΔF/F traces from one ROI each from apical tuft, apical trunk and proximal trunk of a PT neuron in a *Scn2a^+/+^* control mouse. Right, magnified view of the boxed area. **(F)** Similar to (E) for a *Scn2a^+/-^* mouse. **(G – I)** Similar to (D – F) for a frontal cortical IT neuron. ACAd, anterior cingulate area dorsal part. MOs, secondary motor cortex. CP, caudate putamen.

We imaged spontaneous calcium transients from apical dendritic compartments in the awake, head-fixed mouse. We used two-photon microscopy, with the addition of an electrically tunable lens (ETL) placed above the back aperture of the objective. By applying different electrical currents, the ETL can flex quickly, which allows for a rapid change of the focal plane (35). Prior to time-lapse imaging, we would acquire a z-stack from the pial surface to -500 μm with a step size of 5 μm to identify dendritic segments that belong to the same cell. Then time-lapse imaging commenced, which lasted for 30 minutes. For each neuron, we would capture fluorescence transients at three focal planes, targeting one depth each for apical tuft (-50 to - 150 μm), apical trunk (-250 to -400 μm), and proximal dendrite (<-400 μm) (**Fig. 1D**). The overall frame rate was 7.93 Hz. For analysis, we selected regions of interest (ROIs) corresponding to dendritic segments (see **Methods**). We calculated the fractional change in fluorescence Δ*F/F* for each ROI.

In control animals, we saw that the dendritic calcium transients between different compartments of a frontal cortical PT neuron were generally synchronized (**Fig. 1E**). That is, whenever there was a calcium transient in one compartment (e.g., an ROI in the plane of apical tuft), we would observe a corresponding calcium transient in other compartments (e.g., ROIs for the apical trunk and the proximal trunk). However, there were also cases when transients were observed in one compartment but not another, such as the leftmost calcium event in the proximal trunk in **Fig. 1E**, which was not detectable in the apical trunk or apical tuft. We noted the signal-to-noise was typically worse in the ROIs at the apical tuft compared to the apical trunk and proximal trunk (inset, **Fig. 1E**), which was expected given the few fluorophores within the small volume of a dendritic segment in the apical tuft. Relatedly, the decay kinetics for the calcium transients was noticeably longer in the apical and proximal trunks, also consistent with these compartments having larger volume and thus needed more time for the clearance of calcium after an influx event (36). **Fig. 1F** shows example dendritic calcium transients imaged from a frontal cortical PT neuron in a *Scn2a^+/-^*mouse.

Using the same imaging setup, we also characterized the spontaneous calcium transients in the dendrites of frontal cortical IT neurons in the awake, head-fixed mouse. To target a sparse number of IT neurons, we injected a low titer of AAVrg-hSyn-Cre into the contralateral striatum and AAV-CAG-Flex-GCaMP6f in the medial frontal cortex (**Fig. 1C**). Fluorescence images of fixed coronal sections showed sparse labeling of neurons in both superficial and deep layers of the medial frontal cortex, in agreement with the broader distribution of IT neurons (**Fig. 1C**).

Because IT neurons tended to have cell bodies that lie more superficially, we targeted the three focal planes slightly differently for apical tuft (-50 to -150 μm), apical trunk (-150 to -350 μm), and proximal dendrite (<-350 μm) (**Fig. 1G**). **Fig. 1H** shows dendritic calcium transients imaged from a frontal cortical IT neuron in a control animal. **Fig. 1I** shows an example from a *Scn2a^+/-^* mouse.

### *Scn2a* haploinsufficiency impairs dendritic coupling in PT and IT neurons *in vivo*

Although the calcium transients were mostly synchronized across the different dendritic compartments, we also occasionally observed isolated transients in one compartment only. To quantify the coupling between dendrites, we calculated conditional probabilities. Starting from the Δ*F/F(t)* for each ROI, we used the semi-automated OASIS algorithm to infer the underlying electrical events (37). Then, for instance, to determine the amount of transmission from the apical trunk up to the apical tuft, we would consider each event in the apical trunk and tabulate if there was a corresponding event in the apical tuft occurring within ±3 imaging frames (see **Methods**). In an example drawn from actual data (**Fig. 2A**), there were 20 events detected in the apical trunk including 2 events that lacked a counterpart in the apical tuft, hence the conditional probability was 0.9. We considered pairwise correlation as another metric that could be used to estimate coupling, but correlation coefficient may be prone to errors because the calcium transients in different dendritic compartments have different decay kinetics.

**Fig. 2.**
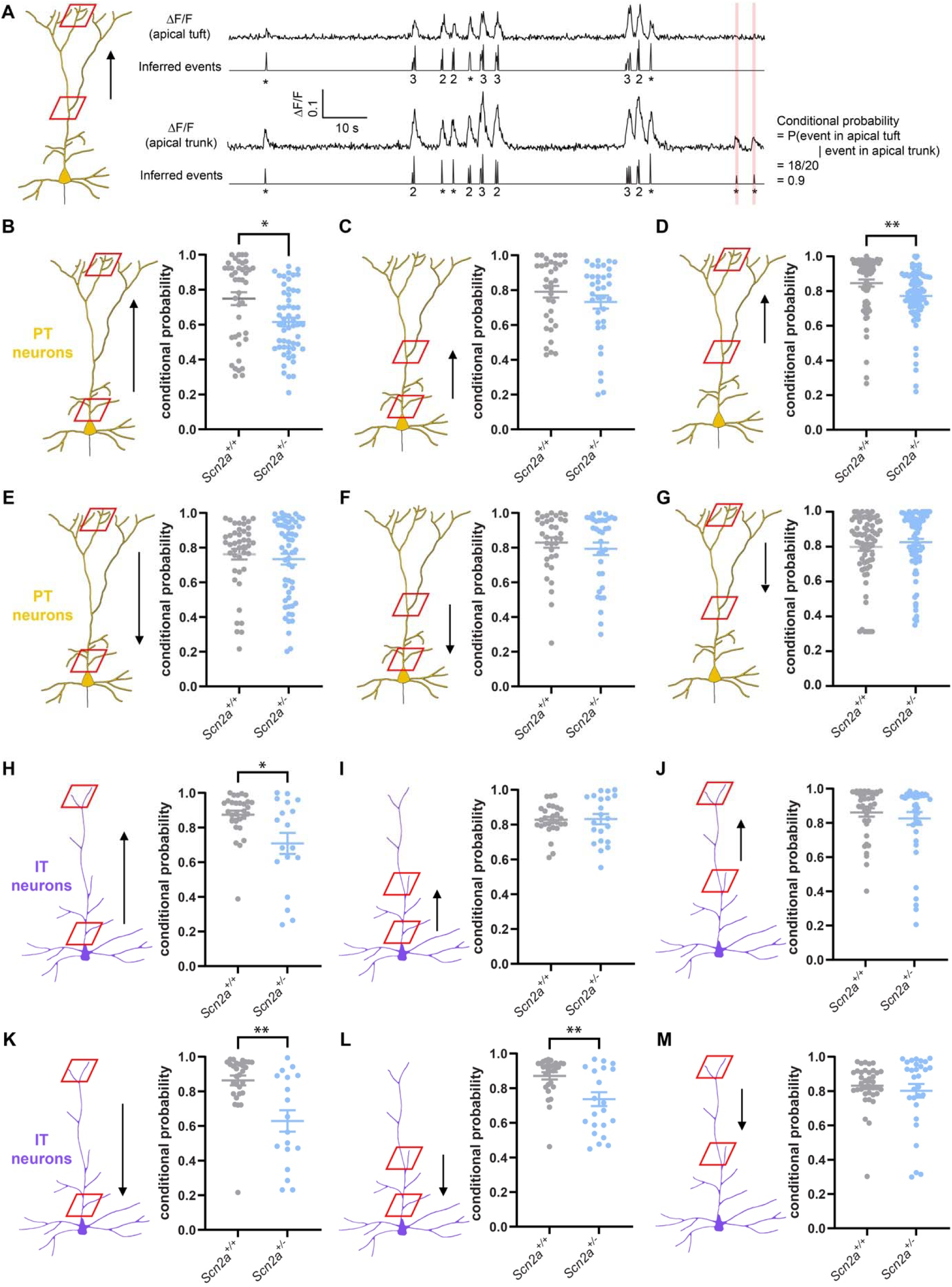
*Scn2a* haploinsufficiency impairs dendritic coupling in PT and IT neurons *in vivo*. Example ΔF/F traces and the corresponding inferred events from the apical tuft and apical trunk of a PT

For PT neurons, we imaged 23 cells in 7 *Scn2a^+/+^* mice (3 males, 4 females) and 31 cells in 6 *Scn2a^+/-^* mice (5 males, 1 female). We calculated conditional probabilities to estimate the coupling between all imaged ROIs (**Fig. 2B – G**). We note that some ROIs could come from the same imaging plane (e.g., several ROIs from the same plane for apical tuft of a neuron), therefore for statistical analysis, we used a mixed effects model, which included random effects terms to account for the hierarchical structure of the data where ROIs come from dendrites that may originate from the same mouse (**Table S1**). In the backward direction, the coupling between the proximal trunk and apical tuft was significantly reduced in *Scn2a^+/-^* mice compared to controls (*Scn2a^+/+^*: 0.75±0.04; *Scn2a^+/-^*: 0.61±0.02; main effect of genotype: *P* = 0.015, mixed effects model; **Fig. 2B**). The other significant effect associated with *Scn2a* haploinsufficiency was reduced coupling between the apical trunk and the apical tuft (*Scn2a^+/+^*: 0.85±0.02; *Scn2a^+/-^* mice: 0.77±0.02; main effect of genotype: *P* = 0.005, mixed effects model; **Fig. 2D**). There was no detected difference in the forward direction for dendritic integration in frontal cortical PT neurons between *Scn2a^+/-^* and control mice (**Fig. 2E – G**).

For IT neurons, we imaged 12 cells in 7 *Scn2a^+/+^* mice (4 males, 3 females) and 13 cells in 5 *Scn2a^+/-^* mice (1 male, 4 females). We found reduced coupling between dendritic compartments in both forward and backward directions (**Fig. 2H – M**). Similar to PT neurons, we found a significant deficit in the coupling between the proximal trunk and apical tuft in the backward direction (*Scn2a^+/+^*: 0.87±0.02; *Scn2a^+/-^*: 0.71±0.06; main effect of genotype: *P* = 0.014, mixed effects model; **Fig. 2H**). For the forward direction, propagation of dendritic signal was impaired from the apical tuft to the proximal trunk (*Scn2a^+/+^*: 0.86±0.03; *Scn2a^+/-^*: 0.63±0.06; main effect of genotype: *P* = 0.002, mixed effects model; **Fig. 2K**), and from the apical trunk to the proximal trunk (*Scn2a^+/+^*: 0.90±0.02; *Scn2a^+/-^*: 0.74±0.04; main effect of genotype: *P* = 0.001, mixed effects model; **Fig. 2L**). Altogether, these results showed that, in many cases, the coupling between dendritic compartments reduced with the heterozygous loss of Na_V_1.2. The coupling was most adversely impacted between the apical tuft and the proximal trunk, which span the greatest distance across the PT and IT neurons in our analysis.

From the events inferred from Δ*F/F* for each ROI, we could determine the rate of the dendritic calcium transients. We found that there were compartment and cell type specific differences in the dendritic calcium transients (genotype × compartment interaction: *P* = 0.021, compartment × cell type interaction: *P* = 0.0003, mixed effects model; **Table S1**). For PT neurons, their dendrites were overall more active in *Scn2a^+/-^* mice than control animals, irrespective of the dendritic compartment (*P* = 5 x 10^-7^, Bonferroni *post hoc* test; **Fig. S1A**). For frontal cortical IT neurons, there was no main difference across genotypes (*P* = 0.8, Bonferroni *post hoc* test), but the frequency of calcium events varied across the dendritic compartments (**Fig. S1D**). Namely, we found that in *Scn2a^+/-^*animals, the calcium event rate was higher in the apical tuft and apical trunk than the proximal trunk (apical tuft vs. proximal trunk: *P* = 0.033, apical trunk vs. proximal trunk: *P* = 0.035, apical tuft vs. apical trunk: *P* = 0.6, Bonferroni *post hoc* test). The reduced event rate in the proximal trunk is consistent with the deficient forward transmission in IT neurons in *Scn2a^+/-^* animals (**Fig. 2K, L**). In addition to event rate, the partial loss of *Scn2a* also affected the amplitude and decay time of the fluorescence transients (**Fig. S1B-C, E-F**; **Table S1**). This analysis demonstrates an aberrant elevation in dendritic calcium events specifically in the PT neuron subpopulation because of *Scn2a* haploinsufficiency, which may compensate for the deficient coupling across dendritic compartments.

### *Scn2a*-haploinsufficient mice were inflexible against changing behavioral strategies

So far, our results have revealed how *Scn2a* haploinsufficiency affects *in vivo* spontaneous activity in the dendrites, but we also want to know the potential alterations during behavior. The dorsal medial frontal cortex in rodents is involved in adaptive decisions involving behavioral flexibility (38). We therefore tested the animals on various decision-making paradigms. Initially, we trained *Scn2a*^+/-^ mice to perform a probabilistic reward reversal task, but observed no alteration in performance compared to control animals (39). Next, we trained *Scn2a*^+/-^ mice to play a standard version of the matching pennies game that we previously adapted for head-fixed mice (40, 41). Matching pennies is a classic two-playing competitive game (42). In this standard version, the two players are the mouse and a computer, with the computer programmed to always play with the same strategy. We again found that *Scn2a*^+/-^ mice performed to a level that was not distinguishable from control animals (**Fig. S2**). We surmised that these tasks may not sufficiently assess flexible behavior, because the animal can rely on the same behavioral strategy for the entire task. For the probabilistic reward reversal, mice can excel by using a finite-state approach, e.g. choosing the high-reward option repeatedly, until encountering a few reward omissions, then switching to the other option (43, 44). For matching pennies, since the computer always used the same strategy, animals could counter by employing a strategy of randomly choosing between the two options (40, 45). Instead, ASD-related cognitive deficits may be accentuated in tasks that encourage flexibility in behavioral strategies.

We therefore designed a new task, still based on matching pennies, but the animal face a computer opponent that can switch strategies. In other words, the rules of the game – payoff matrix and trial timing – always remain the same, but the innovation is that we programmed three different strategies for the computer opponent. Briefly, a head-fixed mouse would play against a computer opponent (**Fig. 3A**). Each trial, the mouse would indicate a choice between the left and right options using a directional tongue lick. The computer would also choose one of the options. The payoff matrix is shown in **Fig. 3B**. If the mouse and computer chose the same option (i.e., both picked left or both picked right), then the mouse would receive a reward of ∼4 μL of water. If the mouse and computer chose different options (i.e., mouse picked left and computer picked right, or vice versa), then the mouse would have no reward. A trial began when a motor actuator advanced the two lick spouts to just in front of the mouse’s head (**Fig. 3C**).

**Fig. 3.**
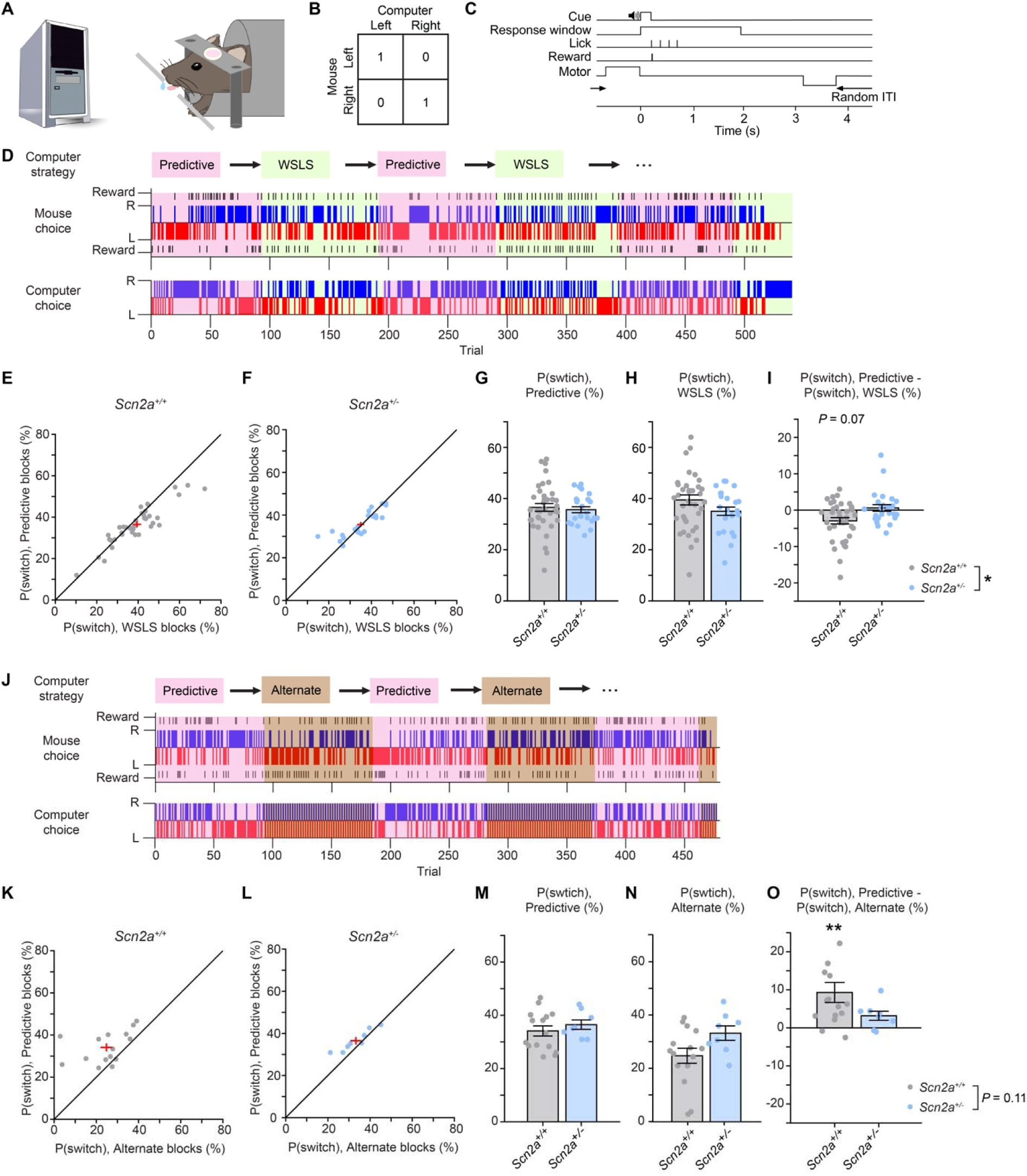
*Scn2a*-haploinsufficient mice were inflexible against changing behavioral strategies. **(A)** Behavioral setup. **(B)** The payoff matrix of the matching pennies game. 1 denotes a reward for the mouse, and 0 denotes no reward. **(C)** The trial timing. An example session when a mouse played matching pennies with a computer opponent that changed strategies between predictive (pink shading) and WSLS (green shading). Top row, the strategy used by the computer. Middle row, the mouse’s choices and outcomes. Bottom row: computer’s choices. Black line, reward.

Then a sound cue was played, which was followed by a 2-s response window. Depending on the mouse’s choice (first lick during the response window) and the payoff matrix, water would be delivered immediately if the mouse won the trial. Subsequently, the motor actuator would retract and there was a random inter-trial interval between each trial. The computer can play with one of three possible strategies. The “predictive” strategy is a strong player that would play by predicting the mouse’s next choice based on the history of past choices and outcomes in that session (same as the strategy used in ref. (40), algorithm 2 in ref. (46), and competitor 1 in ref. (45); see **Methods**). The “WSLS” strategy stands for win-stay, lose-switch, where the computer would pick the same option if it won the last trial or switch options if it lost the last trial. Finally, the “alternate” strategy is when the computer would choose left, right, left, right, and so on, regardless of the mouse’s previous choices and outcomes.

We trained *Scn2a^+/-^* and control mice to play the matching pennies game against a computer opponent using the predictive strategy until the animals became proficient (>35% reward rate for 3 consecutive days; see **Methods and Materials**). Then we tested them on the matching pennies game with the exact same rules, but now against a computer opponent that switched its strategy every 90 – 110 trials during a session. **Fig. 3D** shows a control mouse playing matching pennies when the computer opponent started with the predictive strategy, switched to WSLS, and then toggled for the remainder of the session. We characterized the animals’ performance in 130 sessions from 38 *Scn2a^+/+^*mice (16 males, 22 females) and 108 sessions from 25 *Scn2a^+/-^* mice (15 males, 10 females). The control animals had a probability of switching their option at 36.5±1.5% during predictive blocks and 39.5±2.0% during WSLS blocks (**Fig. 3E**). The *Scn2a^+/-^*animals had a probability of switching their option at 35.6±1.2% during predictive blocks and 35.0±1.1% during WSLS blocks (**Fig. 3F**). Although the differences in performance on aggregate was obscured by variability across individuals (**Fig. 3G, H**), when we specifically analyzed how each animal adapted by calculating its switching probability during predictive block relative to during WSLS block, we detected a significant impairment for the *Scn2a^+/-^* mice to adjust to the changing strategy (*Scn2a^+/+^*: -2.9±0.9%; *Scn2a^+/-^*: 0.6±0.9%; *P* = 0.032, Wilcoxon rank sum test; **Fig. 3I, S3A-F**). The majority of control animals switched more during WSLS blocks than predictive blocks, although this was a trend and not significant (*P* = 0.07, sign test), unlike *Scn2a^+/-^* animals (*P* = 1, sign test). In agreement, logistic regression analysis showed that control animals had a significant tendency to switch following a rewarded choice during WSLS blocks (**Fig. S4**). This is a beneficial adaptation for the control animals, because a player should switch more during WSLS. In fact, to maximize reward rate, an optimal player should be switching half the time against the predictive strategy but switch all the time against the WSLS strategy.

We also trained animals playing matching pennies when the computer opponent was swapping between the predictive and alternate strategies (**Fig. 3J**). We had a smaller data set, which included 60 sessions from 15 *Scn2a^+/+^*mice (7 males, 8 females) and 38 sessions from 8 *Scn2a^+/-^* mice (4 males, 4 females). The control animals had a probability of switching their option at 34.1±1.9% during predictive blocks and 24.8±2.8% during alternate blocks (**Fig. 3K**). The *Scn2a^+/-^* animals had a probability of switching their option at 36.5±1.8% during predictive blocks and 33.3±2.7% during alternate blocks (**Fig. 3L**). In this case, when we quantified how each animal adapts by calculating its switching probability during the predictive blocks relative to during the alternate blocks, we did not detect a significant difference between the genotype (*Scn2a^+/+^*: 9.3±2.6; *Scn2a^+/-^*: 3.2±1.2; *P* = 0.11, Wilcoxon rank sum test; **Fig. 3M-O, S3G-L**), likely owing to the small sample size. Nevertheless, we note that the control animals switched less during alternate blocks than predictive blocks (*P* = 0.007, sign test), whereas the *Scn2a^+/-^* animals again did not adapt (*P* = 0.3, sign test). We speculate that this shift to perseverative choice behavior for control mice was due to their desire to minimize effort, which may relate to a previous study reporting that when the computer’s choice is not contingent on the animal’s decision, the animal would stick to the same option repeatedly (46).

Altogether, these results demonstrate that control mice can adjust their behavior while playing matching pennies with computer opponents employing different strategies. Notably, they switched more frequently against the WSLS strategy but less against the alternate strategy, showing behavioral flexibility in both directions from their likely over-trained responses to the predictive strategy. *Scn2a^+/-^* mice, by contrast, showed significantly reduced adaptability, and did not deviate from their initial trained behavior.

### Task-related signals in the apical dendritic tuft of PT and IT neurons

We observed reduced dendritic coupling during spontaneous activity in both PT and IT neurons in *Scn2a^+/-^*mice, however still unclear is if there may be potential deficit in dendritic signaling during behavior. We therefore hypothesize that dendritic dysfunction in the medial frontal cortex may relate to the inflexibility in decision-making in *Scn2a^+/-^* mice. To investigate this possibility, we imaged dendritic calcium signals from PT and IT neurons while the mouse was playing the matching pennies game against a computer opponent switching between the predictive and WSLS strategies (**Fig. 4A**). The dual viral injection strategy used for **Fig. 1** enabled imaging of the same dendritic tree from single neurons, however the sparse labeling approach had a low success rate, often leading to zero or dense labeling. Here, to increase the yield because we also needed to invest time to train animals, we leveraged the Cre-driver lines *Fezf2-2A-CreER* to target cortical PT neurons and *PlexinD1-2A-CreER* to target cortical IT neurons (47). We injected AAV-CAG-Flex-GCaMP6f into the ACAd and medial MOs portion of the medial frontal cortex of *Fezf2-CreER^+/-^::Scn2a^+/-^*mice, with *Fezf2-CreER^+/-^::Scn2a^+/+^* as controls, or *PlexinD1- CreER^+/-^::Scn2a^+/-^* mice, with *PlexinD1-CreER^+/-^::Scn2a^+/+^* as controls (**Fig. 4B**). We administered tamoxifen to induce Cre-mediated expression of GCaMP6f. We focused on imaging dendritic segments from the apical tuft located 50 – 100 μm below the pial surface (**Fig. 4C, D**).

**Fig. 4.**
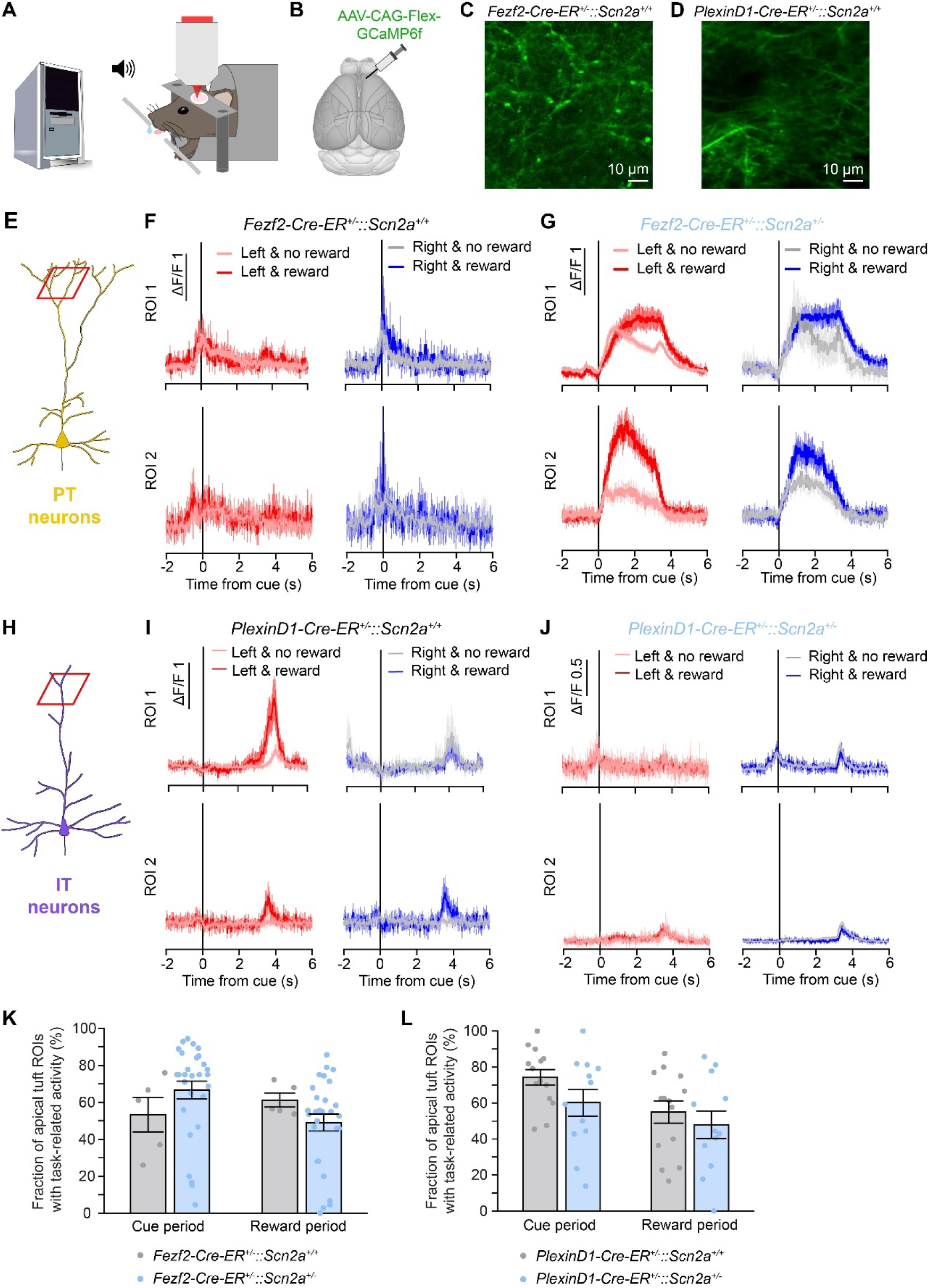
Task-related signals in the apical dendritic tuft of PT and IT neurons. (A) Experimental setup. Viral strategy to express GCaMP6f in PT or IT neurons in the frontal cortex in a mouse.

We aligned fluorescence transients to the time of the sound cue onset, which revealed task-related signals in the apical dendritic tuft in PT and IT neurons (**Fig. 4E-J**). Mostly, fluorescence transients occurred at approximately -1, 0, or 3 s relative to cue onset, corresponding to the times during a trial when the motorized actuator was advancing the lick spouts, the animal was making a lick choice and receiving the outcome, or when the motorized actuator was retracting the lick spouts. **Fig. 4F** shows two ROIs from PT neurons in a control mouse that had task-related activity, although no differential modulation by choice or outcome. By contrast, some apical dendritic tufts of PT neurons in *Fezf2-CreER^+/-^::Scn2a^+/-^*mice exhibited outcome selectivity, for example here both ROIs 1 and 2 larger Δ*F/F* in rewarded trials relative to unrewarded trials (**Fig. 4G**). **Fig.4I** shows example ROIs from IT neurons in a control animal.

Fluorescence increased in ROI 1 when the animal chose left and received a reward, relative to all other conditions. This indicates a preference for the choice *×* outcome interaction, specifically for left and reward. ROI 2 exhibited higher fluorescence when there was a reward together with either a left or right choice, which reflected an outcome-related signal. **Fig. 4J** shows two example ROIs from a control *PlexinD1-CreER^+/-^::Scn2a^+/-^* mouse. These were task-related signals but not selective for specific choice or outcome.

In total, for PT neurons, we imaged 129 ROIs during 5 sessions from 2 *Fezf2-CreER^+/-^::Scn2a^+/+^*mice (2 males) and 861 ROIs during 30 sessions from 8 *Fezf2-CreER^+/-^::Scn2a^+/-^* mice (5 males, 3 females). To quantify the fraction of ROIs that exhibited task-related signals, we determined whether there was a significant difference in Δ*F/F* between the cue period (t = -0.5 to 0.5 s) and baseline (t = -2 to -1 s) or between the outcome period (t = 1 to 2 s) and baseline (t = -2 to -1 s). The majority of ROIs exhibited task-related signals, and the proportions were not different with the partial loss of *Scn2a* than controls for cue period (t = -0.5 – 0.5 s; control: 53.4±9.4%; *Fezf2- CreER^+/-^::Scn2a^+/-^*: 66.7±4.8%; *P* =0.2, Wilcoxon rank sum test) or outcome period (t = 1 – 2 s; control: 61.2±3.7%; *Fezf2-CreER^+/-^::Scn2a^+/-^*: 49.1±4.6%; *P* =0.3, Wilcoxon rank sum test; **Fig. 4K**). For IT neurons, we imaged 357 ROIs from IT neurons during 17 sessions from 4 *PlexinD1- CreER^+/-^::Scn2a^+/+^* mice (2 males, 2 females) and 271 ROIs during 14 sessions from 4 *PlexinD1-CreER^+/-^::Scn2a^+/-^*mice (4 males). Similarly, most ROIs had task-related signals, with the proportions not different across genotype for cue period (control: 74.3±4.3%; *PlexinD1- CreER^+/-^::Scn2a^+/-^*: 60.2±7.5%; *P* = 0.1, Wilcoxon rank sum test) or outcome period (control: 55.0±6.2%; *PlexinD1-CreER^+/-^::Scn2a^+/-^*: 47.8±7.7%; *P* =0.6, Wilcoxon rank sum test; **Fig. 4L**). This analysis indicates that most dendritic segments of frontal cortical PT and IT neurons had task-related calcium transients while the mice were engaged in the matching pennies task.

### Reward- and strategy-related signals were altered in the dendrites of *Scn2a*- haploinsufficient mice

To characterize the factors that account for the calcium signals in the apical dendritic tufts of PT and IT neurons during matching pennies, we used multiple linear regression. For each ROI, we fitted a regression model to determine how the fluorescence transient Δ*F/F*(*t*) relates to the choices (left or right), outcomes (reward or no reward), and their interactions for the current and past trials, as well as strategies (predictive or WSLS) and their interactions (**Fig. 5A**). For PT neurons, the regression analysis showed that fluorescence transients did not significantly encode any of the task variables tested in control animals (**Fig. 5B, C**). Surprisingly, with the partial loss of *Scn2a* in the in *Fezf2-CreER^+/-^::Scn2a^+/-^*mice, we observed that some dendritic calcium signals began to display preference for specific outcome of the current trial and the strategy (**Fig. 5D**). By contrast, when we applied the same multiple linear regression for ROIs from IT neurons, the analysis revealed that ∼7% of the ROIs were modulated by the outcome of the current trial in control animals (**Fig. 5E, F**). The strategy of the computer opponent was robustly represented, especially around the time of cue when decisions were made. These outcome- and strategy-selective signals were absent in apical dendritic tufts in the *PlexinD1- CreER^+/-^::Scn2a^+/-^* mice (**Fig. 5G**). To summarize, unlike earlier studies that reported reward-related firing changes persisting for a few trials in a substantial portion of frontal cortical neurons (38, 48–52), here we observed that outcome- and strategy-related signals were more transient and only present in a small subset of apical dendritic tufts. The fraction of ROIs that has task-related representations was low, which could be due to the low signal-to-noise ratio in our dendritic imaging conditions, or due to substantial contributions from uninstructed movements not associated with the task (53, 54) or arousal-related factors (55), which were not captured by our model. Despite this caveat, our results suggest that *Scn2a* haploinsufficiency alter the typical pattern of dendritic signals in the apical tuft of frontal cortical PT and IT neurons during decision-making.

**Fig. 5.**
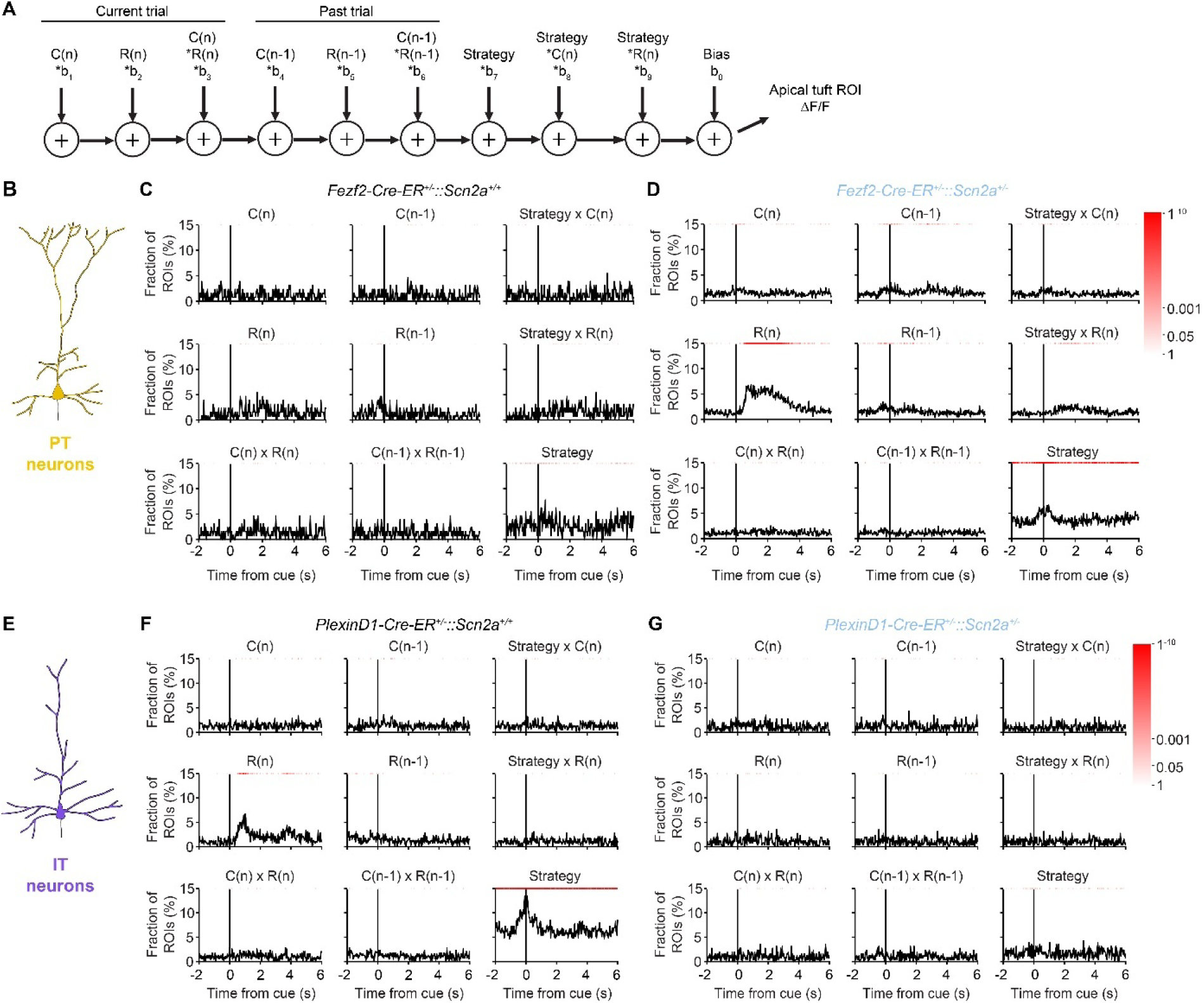
Reward- and strategy-related signals in the dendrites were altered in *Scn2a*-haploinsufficient mice. **(A)** The multiple linear regression model used to fit the ΔF/F of a ROI from the apical tuft of PT and IT neurons. **(B)** Diagram of a PT neuron. **(C)** Fraction of ROIs in *Fezf2-CreER^+/-^::Scn2a^+/+^* control animals with statistically significant (*P <* 0.01) regression coefficients for the various predictor variables. Red shading indicates *P*-value calculated based on the binomial test. **(D)** Similar to **(C)** in *Fezf2-CreER^+/-^::Scn2a^+/-^* mice. **(E)** Diagram of an IT neurons. **(F)** Similar to **(C)** in *PlexinD1-CreER^+/-^::Scn2a^+/+^* mice. Similar to **(C)** in *PlexinD1-CreER^+/-^::Scn2a^+/-^* mice.

## DISCUSSION

The results in this study support three main conclusions. One, *Scn2a* haploinsufficiency impairs the coupling between apical and proximal dendritic compartments in PT and IT pyramidal neurons in the medial frontal cortex. Two, *Scn2a^+/-^*mice exhibited cognitive rigidity, where they were unable to adapt to the changing strategy of a computer opponent in a matching pennies game. Three, reward- and strategy-related signals in the dendrites of frontal cortical PT and IT neurons were disrupted in *Scn2a*-haploinsufficient mice, hinting at a dendritic substrate for the inflexible behavior during decision-making.

Numerous studies have investigated the coupling between dendritic and somatic compartments in cortical pyramidal neurons *in vivo*. In anesthetized animals, layer 5 pyramidal neurons in the barrel cortex exhibit varied coupling between the apical dendrites and proximal soma (56).

Specifically, large calcium signals in the apical dendrites are associated with burst firing in the soma. These correlated events may be elicited by whisker stimulation or by backpropagating action potentials. The fluorescence transients detected in our study are likely associated with these burst firing events rather than single action potentials, as indicated by the relatively low rate of calcium events and the sensitivity of the genetically encoded calcium indicator under our imaging conditions (36). Our observation of high level of dendritic coupling in control animals also aligns with other prior works. A previous study suggests that layer 5 pyramidal neurons in the anterior cingulate cortex may exhibit more reliable signal propagations between soma and dendrite compared to those in the sensory cortex (57). In the awake animal, increased background excitatory activity would elevate dendritic excitability, which should strengthen the coupling between somatic, proximal, and apical compartments (58, 59). Indeed, other recent studies in awake mice reported a close correspondence of calcium transients between distal apical dendrites and locations proximal to the cell body (22, 27, 60). Our results are in agreement with these earlier studies to show a high degree of dendritic coupling in the medial frontal cortex of awake animals.

*Scn2a* encodes the voltage-gated sodium channel Na_v_1.2, which localizes to dendrites of neocortical pyramidal cells after the first postnatal week, and is expected to support dendritic excitability. Indeed, in brain slices, one could evoke dendritic calcium signal via backpropagation by stimulating the cell body, and this signal was reduced with the heterozygous loss of Na_v_1.2 (15). Our study adds to this finding by showing the impact of *Scn2a* haploinsufficiency on the transmission of signals in dendrites *in vivo* in awake animals. We further found cell-type specific differences. PT neurons had impaired coupling affecting transmission from the proximal and apical trunk up to the apical tuft in *Scn2a^+/-^* animals. Accompanying this decoupling in the backward direction is an aberrant elevation in the number of calcium events in the apical tuft. By contrast, IT neurons had diminished dendritic signaling in both the forward and backward directions, and its calcium event rates were not changed by *Scn2a* haploinsufficiency. Though we note that a caveat of the current study is that we are estimating directional effects based on conditional probability, but have not directly measured the propagation of calcium signals which would require much higher temporal resolution for the imaging. The aberrant calcium event rates may relate to the paradoxical increase in action potential excitability seen in cortical pyramidal neurons following the deletion of *Scn2a* (61, 62). For future studies, knowing the distribution of Na_v_1.2 in the dendritic tree of individual frontal cortical PT and IT neurons would allow us to further interpret and model and current data.

Our motivation for selecting a decision-making task was based on two main considerations. First, individuals with ASD exhibit core deficits in social interaction and impairments in decision-making, particularly in computations related to mentalizing and theory of mind (63, 64). Because complex social behaviors are challenging to study directly, we sought tasks that incorporate social and strategic elements as a proxy. One effective approach to probing these deficits in both humans and animal models is through competitive and cooperative games (42, 65, 66).

Second, we aimed to use a task that could be implemented in head-fixed mice, allowing us to perform subcellular-resolution imaging of dendritic signals during behavior. For instance, while attentional set-shifting tasks are well established for assessing cognitive flexibility and have been successfully used in ASD mouse models (67), most implementations involve freely moving mice using food bowls, operant nose pokes, or touchscreens (but see (68), which may be a future direction). These considerations led us to test the *Scn2a* mouse model using matching pennies, which is a classic competitive game paradigm. This approach is relatively untested in rodents, but builds on more extensive previous human studies that linked autism-related traits to performance in strategic games such as the stag-hunt (69), repeated rock-paper-scissors (70), and the dictator game (71). Interestingly, some studies suggest that individuals with ASD may not always have deficits but may instead show enhanced rationality and even outperform controls in certain game contexts (72), highlighting the utility of this approach and nuanced nature of decision-making deficits in autism.

In this study, we observed a subtle but statistically significant behavioral deficit for the *Scn2a^+/-^*mice. There are several reasons why the detected effect was subtle. First, there is individual variability across animals. We tried to minimize variability by carefully standardizing the training environment, however there is still variation in switch rate across individuals of the same genotype. Second, these mice were over-trained. Animals were shaped by first learning matching pennies, where they played repeatedly for tens of thousands of trials against a computer opponent employing the predictive strategy. Once they became proficient, we tested the animals against a computer opponent with changing strategies. Third, likely the strongest factor, computer changed its strategy unbeknownst to the animal. Our analysis was based on dividing trials into blocks based on the computer’s strategy, but the mouse needed time to adapt, thus would still play initially in a new block as if it was playing against the old opponent. For future studies, a more powerful way to assess changes in the mouse’s behavior would be to divide trials into blocks based on the mouse’s latent strategy.

During behavior, reward- and strategy-related signals were diminished in frontal cortical IT neurons of *Scn2a^+/-^* animals. Prior studies have shown that causally silencing neural activity in the frontal cortical region can impair behavioral adjustments (38, 73), therefore our observed loss of task-related signals may underpin the cognitive rigidity. Notably, our results match a prior study identifying reward-related signals in the dendrites of IT, but not PT, pyramidal neurons in the somatosensory cortex in behaving mice (25). Unexpectedly, we observed an aberrant gain of task-selective signals in the dendrites of PT neurons after the partial loss of *Scn2a*, suggesting that the relevant synaptic inputs not be fully lost, but rather maladaptively wired to alter their processing in the medial frontal cortex. Given that frontal cortical PT and IT neurons underpin distinct aspects of motor and decision-related behaviors (74, 75), the cell-type specific deficits provide insights into the circuit mechanisms in ASD.

Despite the strong causal link from *SCN2A* to ASD, prior studies of the heterozygous *Scn2a^+/-^* mice reported only mild behavioral changes. For adult animals, there was no notable change in many behavioral assays designed to test locomotion, anxiety, social interactions, and memory (15, 76). There were some phenotypes for *Scn2a^+/-^* mice, such as decreased ultrasonic vocalization (76), novelty-induced hyperactivity (77), enhanced social interaction (78), and elevated long-term fear memory (77, 78). Phenotypes such as increased responses to lights-on stimuli, daytime hypoactivity, and night-time hyperactivity have also been described in zebrafish models with mutations in the related *Scn1lab* (79). There have been reports of spatial learning deficits in *Scn2a*-haploinsufficient mice (80, 81). A particularly promising trait for translational research is impaired vestibulo-ocular reflex, which is cerebellum-dependent and can be observed in both humans and rodents (82). However, these behavioral phenotypes are difficult to relate to the intellectual deficits observed in humans (12–14). In this study, building on the classic matching pennies game, we developed a task that incentivizes cognitive flexibility.

Testing a large cohort of *Scn2a^+/-^* mice and *Scn2a^+/+^* littermate controls, we demonstrated that control animals could adjust bidirectionally to the shifting behavioral strategies, but mice with *Scn2a* haploinsufficiency failed to adapt. We believe our findings have translational significance, because the same competitive or cooperative game can be tested on animal models and individuals with ASD. In fact, one recent study reported a similar variant of matching pennies, where they programmed computer opponents to play with either a cooperative or competitive strategy and evaluated how human participants would adapt (83). We expect cross-species tasks will facilitate the translation of treatments from animal models to humans.

Understanding how ASD risk genes affect dendritic function is crucial because dendrites play a central role in how neurons receive, integrate, and process information. Here our data provide evidence that the heterozygous loss of the high-confidence risk gene *Scn2a* reduces dendritic coupling in the medial frontal cortex and impairs flexible decision-making. By linking the risk gene to dendritic alterations *in vivo* and during behavior, our results provide mechanistic insights into the cortical circuit deficits underlying the cognitive symptoms observed in ASD, which ultimately may advance diagnostic and treatment strategies for the disorder.

## Acknowledgments

This work was supported by a Simons Foundation Autism Research Initiative Pilot Award (A.C.K.). Additional support was provided by NIH U01NS128660 (A.C.K.), China Scholarship Council-Yale World Scholars Fellowship (H.W.), Cornell Engineering Learning Initiatives (J.I.), NIH training grants T32GM007205 (N.K.S.), NIH fellowship F30MH129085 (N.K.S.), and NIH instrumentation grant S10OD032251 (Cornell Biotechnology Resource Center Imaging Facility).

## Contributions

H.W., L.S., and A.C.K planned the study. H.W. and L.S. performed experiments. J.I. and T.S.J. assisted with the behavioral studies. N.K.S. assisted with imaging experiments. H.W. and L.S. analyzed the data. J.Q. assisted with analysis of the behavioral data. K.J.B. helped with experimental design and discussions. H.W. and A.C.K. drafted the manuscript. All authors reviewed the manuscript before submission.

## Data availability

Data and code associated with the study will be available on https://github.com/Kwan-Lab.

## METHODS

### Animals

*Scn2a^+/-^* mice were originally generated via homologous recombination (84) and sourced from the University of California San Francisco. C57BL/6J (Stock #000664). *Fezf2-2A- CreER* (B6;129S4-*Fezf2^tm1.1(cre/ERT2)Zjh^*/J, Stock #036296) and *PlexinD1-2A-CreER1* (B6;129S4- *Plxnd1^tm1.1(cre/ERT2)Zjh^/J*, Stock #036296) mice (47) were purchased from Jackson Laboratory. A *Scn2a^+/-^* mouse and a C57BL/6J mouse were used as breeding pairs to generate *Scn2a^+/+^* and *Scn2a^+/-^* mice. A *Scn2a^+/-^* mouse and a *Fezf2-2A-CreER^+/+^*mouse were used as breeding pairs to generate *Fezf2-2A-CreER^+/-^::Scn2a^+/+^* and *Fezf2-2A-CreER^+/-^*::*Scn2a^+/-^* mice. A *Scn2a^+/-^* mouse and a *PlexinD1-2A-CreER^+/+^* mouse were used as breeding pairs to generate *PlexinD1- 2A-CreER^+/-^::Scn2a^+/+^* and *PlexinD1-2A-CreER^+/-^*::*Scn2a^+/-^* mice. Ear punch was used for genotyping, in which the inserted gene *Neo^r^* was used to identify the heterozygous loss of *Scn2a.* Mice were housed in groups of 2 – 4 animals with 12-hr-dark and 12-hr-light cycle control (lights off at 7pm at Yale University; lights off at 8pm at Cornell University). Animal procedures were approved by the Institutional Animal Care and Use Committees at Yale University and Cornell University.

### Surgical Procedures

Prior to surgery, the mouse was injected with dexamethasone (3 mg/kg, i.m.; #024751, Henry Schein Animal Health). Animals were anesthetized with 2 - 3% isoflurane in oxygen for induction before the surgery. The isoflurane was lowered to 1 - 1.5% during the surgical procedures. The mouse was placed on a water-circulating heat pad (#RT-0515, Kent Scientific) in a stereotaxic surgical frame (Model 900, David Kopf Instruments). The body temperature was maintained at 38°C. Petrolatum ophthalmic ointment (#IS4398, Dechra) was applied to cover the eyes of the mouse. The hair of the scalp was removed by scissors. The skin was disinfected using an alcohol prep pad (#191089, McKesson), and pushed aside to expose the skull. The scalp was wiped with ethanol and povidone-iodine. A small craniotomy was made above each of the target regions using a handheld dental drill (#HP4-917, Foredom). A borosilicate glass pipette was inserted into the brain and adeno-associated virus (AAV) was delivered intracranially into the target region using an injector (Nanoject II/III Auto-Nanoliter Injector, Drummond Scientific). The injection was performed with a series of 4.6 nL pulses with 30 s interval between pulses. To reduce backflow of the virus, we waited 5 minutes before retracting the glass pipette. For each target region (i.e., medial frontal cortex, striatum, or pons), four injections were made at sites corresponding to four vertices of a 0.2 mm-wide square centered at the center coordinates of the target region. During the procedure, artificial cerebrospinal fluid (ACSF, 135 mM NaCl, 5 mM HEPES, 5 mM KCl, 1.8 mM CaCl2, 1 mM MgCl2; pH 7.3) was applied to keep the cortical surface moist. After injecting all targeted regions, the craniotomies were covered with silicone elastomer (#0318, Smmoth-On, Inc.), and the skin was sutured (#1265B, Surgical Specialties Corporation). Animal was given carprofen (5 mg/kg, s.c.; #002459, Henry Schein Animal Health) after the surgery and on each of the following 3 days.

For imaging spontaneous dendritic calcium transients in PT neurons, 184 nL of AAVrg-hSyn-Cre-WPRE-hGH (1:200 diluted in phosphate-buffered saline (PBS; #P4417, Sigma-Aldrich)) was injected into the pons (anteroposterior (AP): -3.4 mm, mediolateral (ML): -0.7 mm, dorsoventral (DV): -5.2 mm; relative to bregma) and 147.2 nL of AAV1-CAG-Flex-GCaMP6f- WPRE-SV40 (1:10 diluted in PBS) was injected into the ACAd and medial MOs region of the medial frontal cortex (AP: 1.5 mm, ML: -0.4 mm; DV: -1.0 mm) of *Scn2a^+/-^* mice and *Scn2a^+/+^* littermate control mice. For imaging spontaneous dendritic calcium transients in IT neurons, 184 nL of AAVrg-hSyn-Cre-WPRE-hGH (1:200 diluted in PBS) was injected into the contralateral striatum (AP: 0.6 mm, ML: 2.2 mm, DV: -2.8 mm) and 147.2nL of AAV1-CAG-Flex-GCaMP6f- WPRE-SV40 (1:10 diluted in PBS) was injected into the medial frontal cortex (AP: 1.5 mm, ML: - 0.4mm; DV: -1.0mm) of *Scn2a^+/-^*mice and *Scn2a^+/+^* littermate control mice.

At 3 weeks after the viral injection, there was a second surgery to implant a glass window. The pre- and post-operative steps were the same. An incision was made to remove skin and expose the skull. The skull was cleaned by wiping with the ethanol and povidone-iodine. A 3 mm-diameter circular craniotomy centering at the medial frontal cortex was made using the handheld dental drill. ACSF was applied to keep the dura moist. A window was constructed using two pieces of glass coverslips (3 mm diameter, 0.15 mm thickness; #640720, Warner

Instruments) that were bonded with ultraviolet light-curing optical adhesive (#NOA 61, Norland Products) using an ultraviolet illuminator (#2182210, Loctite). This two-layer glass window was placed into the craniotomy, and super glue adhesive (Henkel Loctite 454) was used to secure the window to the surrounding skull. After 20 minutes when the glue was dry, a stainless steel headplate (eMachineShop; design available at https://github.com/Kwan-Lab/behaivor-rigs) was placed centered on the glass window using the super glue (All-purpose glue, Krazy) and a quick adhesive cement system (Metabond, Parkell). Mouse would recover for two weeks after the window implant prior to imaging experiments.

For imaging dendritic calcium transients during behavior, the procedures were performed in a single surgery. The pre- and post-operative steps were the same. The hair of the scalp was removed by scissors. An incision was made to remove skin and expose the skull. The skull was cleaned by wiping with the ethanol and povidone-iodine. A 3 mm-diameter circular craniotomy centering at the medial frontal cortex was made using the handheld dental drill. ACSF was applied to keep the dura moist. To target PT or IT neurons, 147.2 nL of AAV1-CAG-Flex- GCaMP6f-WPRE-SV40 (1:10 diluted in PBS) was injected into the medial frontal cortex (AP: 1.5 mm, ML: -0.4 mm; DV: -1.0 mm) of *Fezf2-2A-CreER^+/-^::Scn2a^+/-^*, *Fezf2-2A-CreER^+/-^::Scn2a^+/+^*, *PlexinD1-2A-CreER^+/-^::Scn2a^+/-^*, or *PlexinD1-2A-CreER^+/-^::Scn2a^+/+^* mice. The other injection parameters were the same as described in the previous section. A four-layer glass window was made by bonding 1 layer of 4 mm-diameter glass together with 3 layers of 3 mm-diameter glass using the ultraviolet light-curing optical adhesive. The thicker glass window was needed to minimize the motion of the brain, which was more vigorous during behavior than when imaging spontaneous activity. This glass window was placed in the craniotomy and secured by the super glue adhesive. After 20 minutes, a stainless steel headplate was placed centered on the glass window with the super glue and a quick adhesive cement system.

### Tamoxifen injection

Tamoxifen was used to induce Cre-dependent gene expression when using *Fezf2-2A-CreER^+/-^::Scn2a^+/-^*, *Fezf2-2A-CreER^+/-^::Scn2a^+/+^* mice, PlexinD1*-2A-CreER^+/-^ ::Scn2a^+/-^*, and *PlexinD1-2A-CreER^+/-^::Scn2a^+/+^* mice. Tamoxifen (#T4648, Sigma-Aldrich) was dissolved in corn oil (#C8267, Sigma-Aldrich) at a concentration of 20 mg/mL. The solution was placed for 1 hour at 37 °C in an ultrasonic water bath to help dissolve the compound. The solution was aliquoted into 2 mL Eppendorf tubes and stored at -20°C. Prior to injection, the aliquoted tamoxifen was thawed at 4°C. Tamoxifen (75 mg/kg, i.p.) was injected for 5 consecutive days starting from at 3 days after surgery. Animals started water restriction and behavior training at least 7 days after the last tamoxifen injection.

### Behavioral setup

The operant training setup for the head-fixed mouse was based on the design from our prior studies(38, 50). Detailed instruction to construct the apparatus is available at https://github.com/Kwan-Lab/behavioral-rigs. The mouse’s head stabilized via the headplate using a stainless-steel piece (eMachineShop). The mouse sat in an acrylic tube (8486K433; McMaster-Carr), which limited gross movements but allowed postural adjustments. A lick port with two water spouts was positioned in front of the mouse. The spouts were blunted 20-gauge stainless-steel needles. The lick port was placed on a motorized linear actuator (0.8” stroke, 4.5 lb; B07ZJ4B3WW, DC House), which was powered by an L293D motor driver (497-2936-5-ND, DigiKey). The actuator enabled us to move the lick port to a location accessible to the mouse at the start of a trial and then retract the lick port at the end of the response window. When the mouse licked either spout, the contact was recorded by a battery-powered lick detection electronic circuit, which sent signals to the computer via a data acquisition unit (USB-201, Measurement Computing) and logged by the Presentation software (Neurobehavioral System Inc.). Water delivery from each lick spout was controlled by a solenoid fluid valve (MB202-V-A-3- 0-L-204; Gems Sensors & Controls). When there was a reward, ∼4 μL of water was delivered by sending a signal via the data acquisition unit to the water delivery electronic circuit to activate the solenoid. A pair of speakers (S120, Logitech) were placed in front of the mouse to play the sound cue. For training, the entire setup was placed inside a behavior box with walls lined with soundproof acoustic foams (5692T49, McMaster-Carr).

### Behavioral training – matching pennies

All the tasks were programming using the scripting language in the Presentation software. Mice were water restricted for 48 hours before the behavioral training. For the first training stage, mice were habituated to head restraint and the lick spouts were always positioned in front of the mouse (i.e., the actuator did not move). They could lick either lick spout for water until they are satiated. The first stage lasted 2–3 days, with the aim of helping the mouse get used to the head restraint and the licking circuit. For the second training stage, at the start of a trial, the actuator started to advance. We measured that the actuator took 0.85 s to bring the lick spout to the front of the mouse. Therefore, 0.85 s after the start of a trial, a 5 kHz sound cue lasting 0.2 s was played and a 2-s response window began. During the response window, the first lick to either side of lick spouts will lead to the immediate delivery of ∼4 μL of water to the contacted spout and triggered a fixed 3-s period for consumption. If no lick was detected, the fixed 3-s period would be applied at the end of the 2-s response window. At the end of the fixed 3-s period, the actuator started to retract and an intertrial interval was applied until the start of the next trial. The duration of the intertrial interval was drawn randomly for each trial, from a truncated exponential distribution with λ = 1/3 and boundaries of 1 and 5. Once the ITI concluded, the next trial would start. The session continued until either the mouse misses for 10 consecutive trials, or if it made 1000 responses. The second stage typically lasted 2 days, with the aim of helping the mouse get used to the moving actuator. When a mouse could lick at least 100 trials per session, it proceeded to the matching pennies game.

For matching pennies task, the timing of events in a trial was exactly the same as the second stage. The first lick to either water spout during the 2-s response window would be the mouse’s choice (left or right) for that trial. According to the payoff matrix of matching pennies, if the mouse’s choice matched the computer’s choice, ∼4 μL of water will be delivered and triggered the fixed 3-s period. If the mouse’s choice did not match the computer’s choice, there was no reward but still triggered the fixed 3-s period. In this standard version of the matching pennies game, the computer opponent made choices using the ‘predictive’ strategy. This strategy is the same as the strategy used in ref. (40), algorithm 2 in ref. (46), and competitor 1 in ref. (45).

Briefly, the computer had knowledge of the mouse’s choice and outcome history since the start of the session. The computer would calculate 9 conditional probabilities that the mouse would choose left given sequences of the preceding N choices (N = 0 – 4) and N choice-outcome combinations (N = 1 – 4). A binomial test was then used to test each of these conditional probabilities against the null hypothesis that the mouse would choose left with a probability of 0.5. If none of the null hypotheses was rejected, the computer would randomly choose between left or right. If one or more hypotheses were rejected, the computer would choose the side with the statistically significant conditional probability that was furthest away from 0.5.

### Behavioral training – matching pennies against a computer opponent with shifting strategies

Mice were trained to play the standard version of the matching pennies game; when a mouse achieved a reward rate of 35% or greater for 3 consecutive days, it was deemed competent, and we would test the animal against a computer opponent with shifting strategies. In this variant, the computer opponent, which we called Tom (after Tom, the strategist who switches strategy to try to catch the mouse Jerry), can play with 1 of 3 different strategies: predictive, WSLS (win-stay lose-switch), and alternate. Tom would use the same strategy for a block of trials, drawn randomly from a uniform distribution ranging between 90 and 110, and then switch abruptly to the different strategy, without any indication to the mouse. For the predictive strategy, this is the same as described above, except Tom would calculate the conditional probabilities based on the mouse’s choice and outcome history since the start of the current block. For the WSLS strategy, Tom would randomly choose left or right for the first trial. If Tom wins, it will stay on that same option for the subsequent trial. But if Tom loses, it will switch to the other option for the next trial. Tom will continue with this logic until the end of the block.

For the alternate strategy, Tom would randomly choose left or right for the first trial. On the next trial, Tom will always choose the other option, and continue to alternate between the two options until the end of the block. At the start of a session, Tom began with the predictive strategy for the first block, then a second strategy (WSLS or alternate) for the second block, the predictive strategy for the third block, the same second strategy for the fourth block, and so on. Hence, Tom switches either between predictive and WSLS or predictive and alternate. The session continued until either the mouse misses for 10 consecutive trials, or if it made 1000 responses.

### Analysis of behavior data

We included all sessions when the mouse responded for at least 50 trials for each block. For the standard version of matching pennies, we had 139 sessions from 16 *Scn2a^+/+^*animals (7 males, 9 females) and 181 sessions from 21 *Scn2a^+/-^*animals (12 males, 9 females). For the matching pennies against a computer opponent with shifting between predictive and WSLS strategies, we had 130 sessions from 38 *Scn2a^+/+^* mice (16 males, 22 females) and 108 sessions from 25 *Scn2a^+/-^* mice (15 males, 10 females). For the matching pennies against a computer opponent with shifting between predictive and alternate strategies, we had 60 sessions from 15 *Scn2a^+/+^*mice (7 males, 8 females) and 38 sessions from 8 *Scn2a^+/-^* mice (4 males, 4 females). Some of the *Scn2a^+/+^* mice were *Fezf2-2A-CreER^+/-^ ::Scn2a^+/+^* or *PlexinD1-2A-CreER^+/-^::Scn2a^+/+^* animals, and some of the *Scn2a^+/-^* mice were *Fezf2-2A-CreER^+/-^::Scn2a^+/-^* or *PlexinD1-2A-CreER^+/-^::Scn2a^+/-^* animals, including all sessions where we performed imaging during behavior.

The behavior data, including animal’s choices and outcomes, lick times, and computer’s strategy, were stored by the Presentation software as text log files. The log files were processed with custom-written code in MATLAB. To compute the probability of switch *P*(switch), we would sum the number of trials in which the choice was different than the prior choice (i.e., animal selected right and the last choice was left, or animal selected left and the last choice was right), and divide by the total number of trials. To compute the probability of reward *P*(reward), we would sum the number of trials in which the animal received a water reward (i.e. animal selected right and the computer selected right, or animal selected left and the computer selected left), and divide by the total number of trials. For each session we would calculate *P*(switch) and *P*(reward) separately using trials in which computer strategy was predictive or WSLS, or predictive or alternate.

We performed a logistic regression analysis to determine how past choices and reinforcers influence the current decision. The logistic regression analysis was based on the following equation:

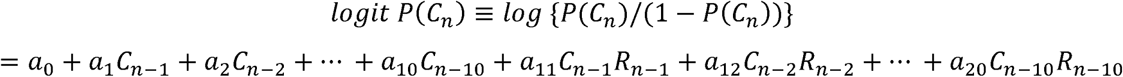

where *P(C_n_)* is the probability of choice in trial *n*, *C_n-1_* to *C_n-10_* are the choices made by the mouse on the previous trials up to 10 trials back, *R_n-1_* to *R_n-10_* are the outcomes received by the mouse on the previous trials up to 10 trials back, and *a_0_* to *a_20_* are the regression coefficients. Choices were coded as -1 for left choice and 1 for right choice. Outcomes were coded as 0 for no reward and 1 for reward. To determine the coefficients, we fit this model to the mouse’s choice behavior by optimizing the negative log-likelihood value. We separately fit two models, either using trials during predictive blocks by concatenating all those trials across all sessions and all mice or using trials during WSLS blocks by concatenating all those trials across all sessions and all mice.

### Two-photon microscopy

For multi-plane imaging of the spontaneous dendritic calcium transients, the experiments were performed with a Movable Objective Microscope (MOM, Sutter Instrument) equipped with a resonant-galvo scanner (Rapid Multi Region Scanner, Vidrio Technologies or Resonant Scan Box, Sutter Instrument). We used a water-immersion 20x objective (XLUMPLFLN, 20X/0.95 N.A., Olympus), with an electrically tunable lens (EL-3-10; Optotune) placed just above the back aperture of the objective. Scanimage 2020 software was used to control the microscope for image acquisition. The excitation source was either a tunable Ti:Sapphire femtosecond laser (Chameleon Ultra II, Coherent) or a fixed-wavelength ultrafast laser (Axon 920-2 TPC, Coherent). The excitation wavelength was 920 nm, and the emission was collected behind a 475 – 550 nm bandpass filter. The intensity of laser was 120 – 180 mW after the objective. Time-lapse images were acquired at a resolution of 512 x 512 pixels. The ETL was set such that 0 V corresponded to the plane of the proximal trunk, and two different voltages were used to rapidly switch between the 3 different focal planes. The voltages related to depth through a linear relationship and was calibrated. The voltages were sent using the fast focus control module in ScanImage. The field of view was slightly larger at the plane of the apical tuft, compared to the plane of the proximal trunk. The frame rate was 7.93 Hz using bidirectional scanning. Initially the mouse was habituated in the two-photon microscope for 3 days. The habituation time increased day by day, from 10 minutes to 40 minutes. Prior to the time-lapse imaging, z-stack images were acquired from the pial surface to -500 μm with a step size of 5 μm. The z-stack images were used to visualize the dendritic tree of a pyramidal neuron, such that we may select specific depths for the time-lapse imaging. For PT neurons, we would choose one depth each within the range for apical tuft (-50 to -150 μm), apical trunk (-250 to -400 μm), and proximal dendrite (<-400 μm). For IT neurons, we would choose one depth each within the range for apical tuft (-50 to -150 μm), apical trunk (-150 to -350 μm), and proximal dendrite (<-350 μm). We would acquire time-lapse images from the three depths for 30 minutes.

For imaging during behavior, we used the same imaging system with the behavioral apparatus set inside the two-photon microscope. There were several other differences. We used a water-immersion 25x objective (N25X-APO-MP, NIKON). The frame rate was 30.03 Hz using bidirectional scanning. The intensity of laser was 100 – 140 mW after the objective. We imaged dendrites in a focal plane that lies between a depth of -50 and -150 μm. To synchronize behavioral and imaging data, a TTL pulse was sent by the Presentation software at the beginning of each trial from the USB-201 board of the behavior system. The imaging system used the TTL pulse as an external trigger to initiate the acquisition of a new imaging file.

For each session, we would initially acquire baseline images for 30 seconds when the animal has not yet started the task. Then we would acquire time-lapse images, triggered by the behavioral setup as the task progressed, until the task was completed.

### Preprocessing of the multi-plane imaging data

For multi-plane imaging of the spontaneous dendritic calcium transients, we made three 10-minute recordings consecutively, which together form the 30-minute session. Each recording generated a multi-page .tiff file saved by ScanImage. The file contains image frames for the three different focal planes interleaved. We would extract the image frames corresponding to each focal plane, and then concatenate to produce the time-lapse images for a focal plane spanning 30 minutes.

We processed with NoRMCorre in MATLAB to correct for non-rigid translational motion. Next, regions of interest (ROIs) were selected from the motion-corrected time-lapse images, by comparing with the initial z-stack images of the individual neuron. The software to draw ROIs was a GUI previously written in the lab. The criteria for ROI selection were as follows: first, the dendrite was not overlapping with any other dendrites; second, the dendrite belonged to the target neuron; third, there were event-like transients with rapid onset with amplitude of more than ∼0.1 and exponential decay during the session. To account for the influence of the neuropil, we calculated the fluorescence signal for each ROI as follows:

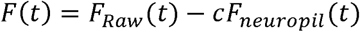

where *F_raw_*(*t*) is the summed pixel value within the ROI at time *t*, *c* is a correction factor set to 0.4, and *F_neuropil_*(*t*) is the summed pixel value in the neuropil area, defined as an annulus shaped area centered at the centroid location of the ROI with inner and outer diameters of 2*r* and 3*r*, where *r* is the radius if we assume the ROI area is a circle. The fractional change in fluorescence Δ*F/F(t)* was calculated as follows:

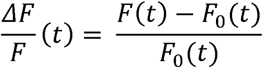

where *F_0_(t)* is the baseline fluorescence, which was the 10^th^ percentile of *F*(*t*) within a 2-min running window centered around time *t*.

### Analysis of the multi-plane imaging data

For PT neurons, we imaged 23 cells in 7 *Scn2a^+/+^* mice (3 males, 3 females) and 31 cells in 6 *Scn2a^+/-^* mice (5 males, 1 female). For IT neurons, we imaged 12 cells in 7 *Scn2a^+/+^* mice (4 males, 3 females) and 13 cells in 5 *Scn2a^+/-^*mice (1 male, 4 females). Details for the data set are provided in **Table S2**. To infer the calcium events from Δ*F/F(t)*, we used an algorithm called Online Active Set method to Infer Spikes, or OASIS (37). To calculate the event rate, we divided the total number of calcium events by the duration of the imaging session.

We estimated the coupling between different dendritic compartments using conditional probability. To calculate conditional probability, for example the probability of event in compartment B given an event in compartment A, we would first identify all events in compartment A. For each event, we examined a window of ±3 imaging frames (∼ ±0.38 s) around its occurrence to check for at least one event in compartment B. If a corresponding event was present, we recorded a 1; otherwise, we recorded a 0. The conditional probability was then calculated by averaging the 1s and 0s across all events in compartment A. As an example, if there were simultaneously detected calcium events in the apical tuft and proximal trunk compartments, then this scenario would count as a 1 for *P*(apical tuft | proximal trunk), and it would also count as a 1 for *P*(proximal trunk | apical tuft). This is reasonable because the temporal resolution of calcium imaging did not allow us to know whether this simultaneously detected event is forward or backward propagating. Insights into the forward- versus backward-direction coupling came from the selective events, such as when there was an event in the apical tuft but not in the proximal trunk. This scenario would count as a 0 for *P*(proximal trunk | apical tuft), but does not count towards anything for *P*(apical tuft | proximal trunk). We emphasize to truly know if an event is forward or backward propagating, we would need to image at a much higher frame rate to capture the temporal sequence of calcium signals. Here conditional probabilities leveraged the case when coupling fails to provide an estimate into the directionality of the coupling.

### Preprocessing of the behavioral imaging data

For imaging during behavior, acquisition of an image file was triggered by a TTL pulse sent by the Presentation software at the start of each trial. Therefore, each trial generated a multi-page .tiff file saved by ScanImage. To analyze the data, we first concatenated the image files in batches of 100 trials to produce several large image files. Then we generated a template for motion correction by calculating the mean projection using the initial baseline images acquired before matching pennies. Based on this template image, we processed all the large image files with NoRMCorre in MATLAB to correct for non-rigid translational motion. Next, regions of interest (ROIs) were selected from the motion-corrected first large image file. The criteria for ROI selection were as follows: first, the dendrite was not overlapping with any other dendrites; second, there were event-like transients with rapid onset with amplitude of more than ∼0.1 and exponential decay. Then, the masks for the ROIs were applied to all other large image files to extract the fluorescence signals. We calculated the fractional change in fluorescence Δ*F/F(t)* using the same steps, except did not correct for the neuropil influence (i.e., c = 0).

### Analysis of the behavioral imaging data

For imaging during behavior, we had 5 sessions from 2 *Fezf2-2A-CreER^+/-^::Scn2a^+/+^* animals (2 males), 30 sessions from 8 *Fezf2-2A-CreER^+/-^ ::Scn2a^+/-^* animals (5 males, 3 females), 17 sessions from 4 *PlexinD1-2A-CreER^+/-^::Scn2a^+/+^* animals (2 males, 2 females), and 14 sessions from 4 *PlexinD1-2A-CreER^+/-^::Scn2a^+/-^* animals (4 males). Details for the data set are provided in **Table S2**. Behavior data were processed with a custom written script in MATLAB. To plot trial-averaged activity, we aligned Δ*F/F(t)* by the time of cue onset, and generated peri-stimulus time histogram by averaging across all trials of the same type (i.e., left & no reward, left & reward, right & no reward, or right & reward). To assess task-related activity, for each ROI, we would compare the trial-by-trial Δ*F/F* between the cue period (t = -0.5 to 0.5 s) and baseline (t = -2 to -1 s) or between the outcome period (t = 1 to 2 s) and baseline (t = -2 to -1 s). To quantify how dendritic calcium transients may relate to choice, outcome and strategy, we constructed a multiple linear regression model:

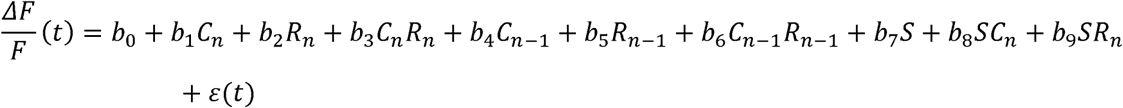

where Δ*F/F(t)* is the fractional change in fluorescence at time *t* relative to cue onset in trial *n*, *C_n_* and *C_n-1_* are the choices made by the mouse on the current trial and the previous trial, *R_n_* and *R_n-1_* are the outcomes received by the mouse on the current trial and the previous trial, *S* is the computer strategy, *b_0_, b_1_,…, b_9_* are the regression coefficients and E(t) is the error variable.

Choices were coded as -1 for left choice and 1 for right choice. Outcomes were coded as 0 for no reward and 1 for reward. Computer strategy was coded as 1 for predictive and -1 for WSLS. The regression coefficients were calculated by fitting the model using the MATLAB function *fitlm* for each imaging session. We fitted for the time period from -2 s before the cue onset to 6 s after the cue onset, with step size of 33.3 ms that matched the imaging frame rate. To collate the results across all sessions, for each predicting variable, we calculated the fraction of dendritic ROIs in which the regression coefficient was significantly different from zero (*P <* 0.01).

### Histology

After the completion of imaging and behavioral studies, mice were perfused with ice-cold PBS and paraformaldehyde solution (PFA, 4% v/v in PBS). The brains were extracted and fixed in 4% PFA at 4°C. The brain was cut into coronal sections at a thickness of 50 μm using a vibratome (#VT1000S, Leica). The brain sections were mounted on slides with glass coverslips. If the brains were to be stored beyond 24 hours before sectioning, PFA was replaced with PBS. For fluorescence imaging of the sections, a confocal microscope (LSM880, Zeiss) equipped with a 10x/0.45 N.A. water objective was used. Images were acquired at 512 x 512 pixels.

### Statistics

All statistics were conducted within MATLAB and R studio. For comparisons across different groups, a linear mixed effects model was implemented with the lme4 package in R. For conditional probability, we constructed a model with fixed effects terms including genotype, compartment pairing, cell type, and all higher order interactions, with ROI per cell per mouse modeled as nested random intercepts. For event rate, amplitude and decay time, we constructed a model with fixed effects terms including genotype, compartment, cell type, and all higher order interactions, with ROI per cell per mouse modeled as nested random intercepts.

When the linear mixed effects model indicated significant main or interaction effects, we performed post hoc pairwise comparison with Bonferroni correction. The specifics for the other statistical tests performed are indicated in the main text and the figure captions.

