## Supplementary Figures for "Autism-associated *Scn2a* haploinsufficiency disrupts *in vivo* dendritic signaling and impairs flexible decision-making"

**Supplementary Figures 1 - 4**

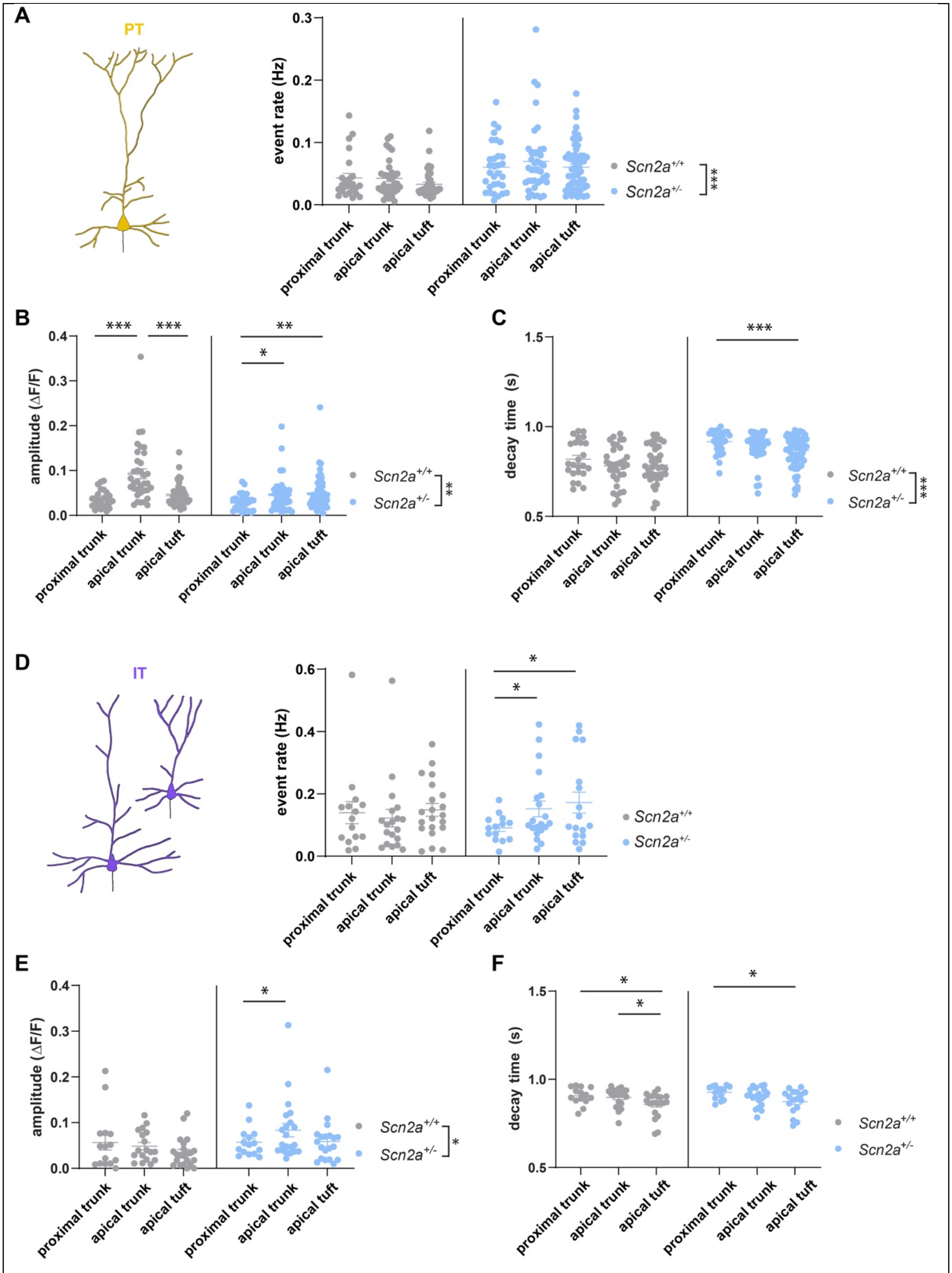


**Fig. S1. Aberrant rates of dendritic calcium events in *Scn2a*$^{+/-}$ mice.**

**(A)** The rate of inferred events for the different dendritic compartments of PT neurons in *Scn2a*$^{+/+}$ control animals and *Scn2a*$^{+/-}$ mice. Each point is a ROI. Mean±SEM.

**(B)** Similar to (A) for the mean amplitude of the inferred events.

**(C)** Similar to (A) for the mean decay time of the inferred events.

**(D - F)** Similar to (A - C) for IT neurons.

*, $P$<0.05. **, $P$<0.01. ***, $P$<0.001. Linear mixed effects model with fixed effects terms of compartment (proximal trunk, apical trunk, or apical tuft), genotype (Scn2a$^{+/+}$ or Scn2a$^{+/-}$), cell type (PT or IT) and all interactions, with ROI per dendrite per mouse modeled as nested random intercepts.

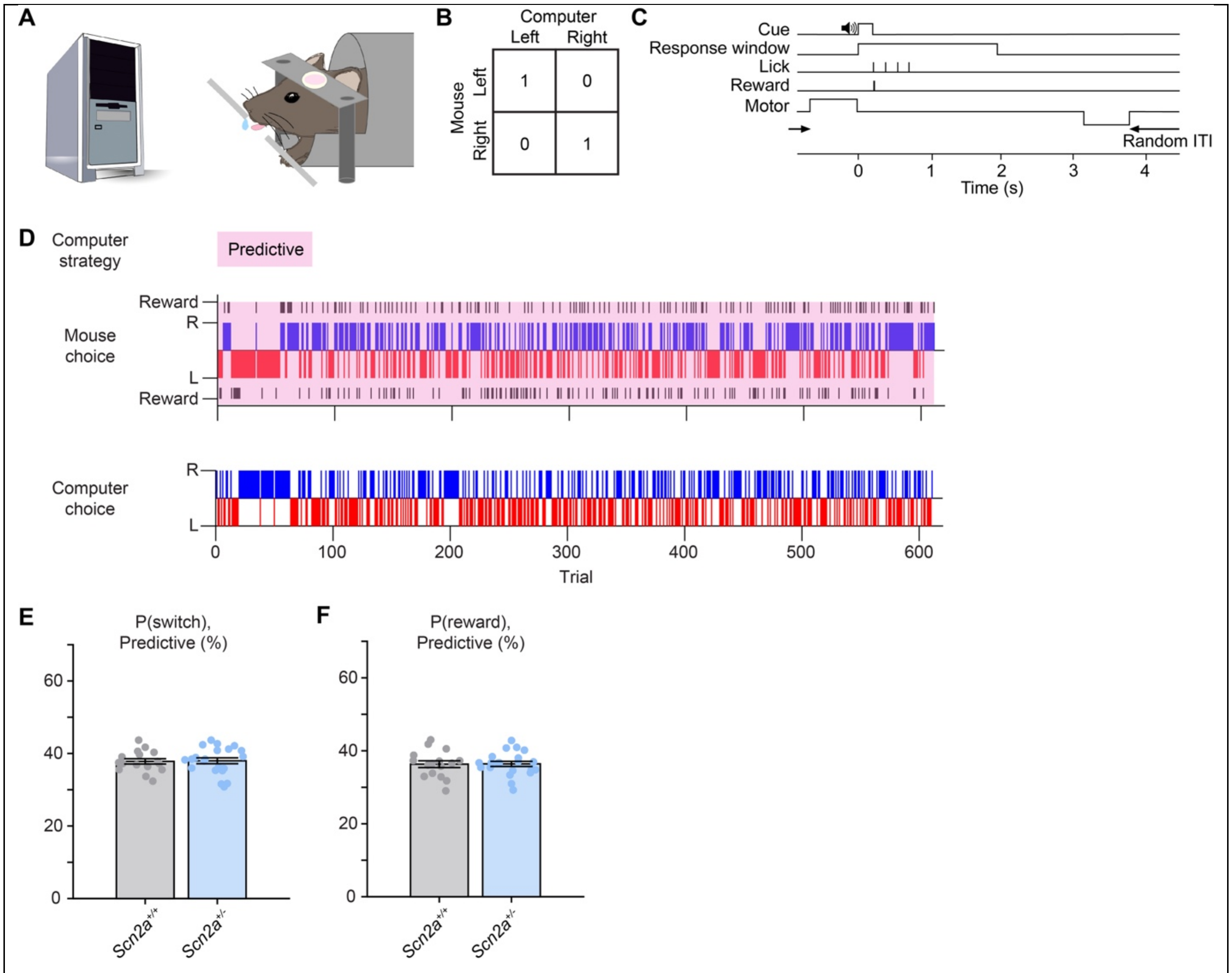


**Fig. S2. *Scn2a*$^{+/-}$ mice play matching pennies against a computer opponent with a fixed strategy.**

**(A)** Behavioral setup.

**(B)** The payoff matrix of the matching pennies game. 1 denotes a reward for the mouse, and 0 denotes no reward.

**(C)** The trial timing.

**(D)** An example session when a mouse played matching pennies with a computer opponent with a fixed "predictive" strategy (pink shading). Top row, the strategy used by the computer. Middle row, the mouse's choices and outcomes. Bottom row: computer's choices. Black line, reward.

**(E)** The switch probability for *Scn2a*$^{+/+}$ control and *Scn2a*$^{+/-}$ mice. Each point represents the mean value of a mouse. Red crosshair, Mean±SEM.

**(F)** Similar to (E) for reward probability**.**

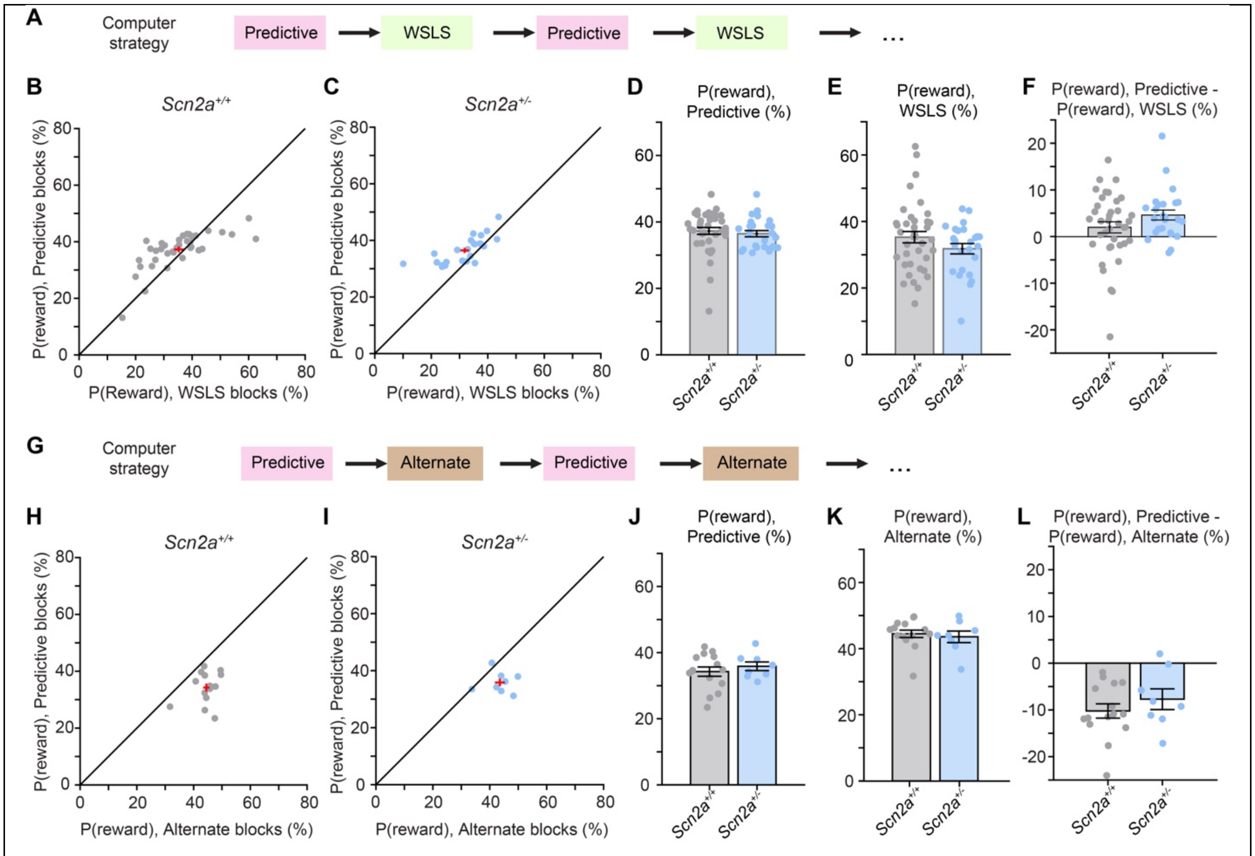


**Fig. S3. Reward rates during matching pennies against computer opponents that changed strategies for *Scn2a*$^{+/-}$ mice and controls.**

**(A)** Mice played matching pennies with a computer opponent that changed strategies between predictive (pink shading) and WSLS (green shading).

**(B)** The reward rate during predictive and WSLS trials for *Scn2a*$^{+/+}$ control mice. Each point represents the mean value of a mouse. Red crosshair, Mean±SEM.

**(C)** Similar to (B) for *Scn2a*$^{+/-}$ mice**.**

**(D)** Bar plot of the reward rate when the computer used predictive strategy. Each point represents the mean value of a mouse. Mean±SEM. Reward rate was 37.3±1.0% for controls and 36.5±0.9% for *Scn2a*$^{+/-}$ mice.

**(E)** Similar to (D) when the computer used WSLS strategy. Reward rate was 35.3±1.7% for controls and 31.8±1.6% for *Scn2a*$^{+/-}$ mice, with the difference not statistically significant ($P$ = 0.24, Wilcoxon rank sum test).

**(F)** Bar plot of the reward rate difference between predictive trials and WSLS trials. Each point represents the mean value of a mouse (*Scn2a*$^{+/+}$: 2.0±1.2%; *Scn2a*$^{+/-}$: 4.6±1.1%; $P$ = 0.27, Wilcoxon rank sum test). Mean±SEM.

**(G) – (L)** Similar to (A) – (F) for a computer opponent that changed strategies between predictive (pink shading) and alternate (brown shading). For predictive trials, reward rate was 34.3±1.4% for controls and 35.9±1.3% for *Scn2a*$^{+/-}$ mice. For alternate trials, reward rate was 44.5±1.1% for controls and 43.6±1.8% for *Scn2a*$^{+/-}$ mice.

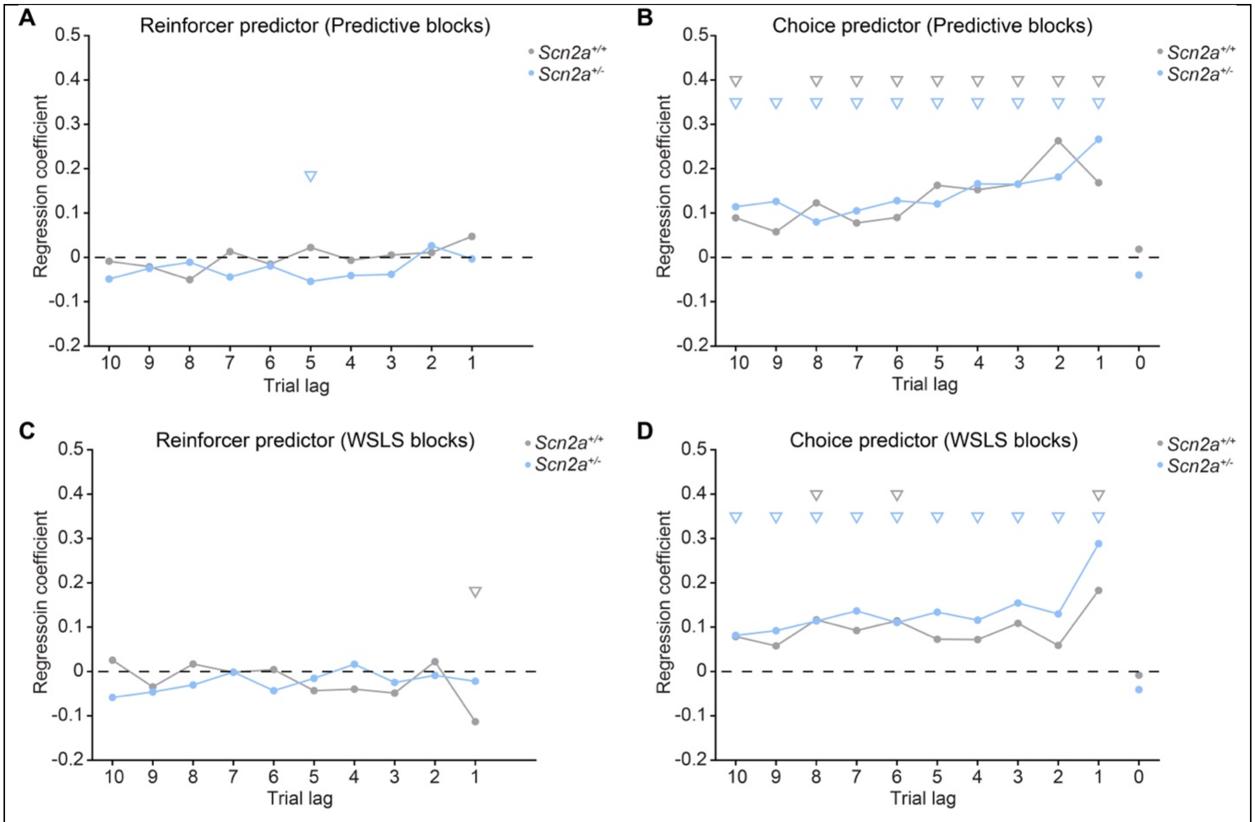


**Fig. S4. Logistic regression analysis of choice behavior for *Scn2a*$^{+/-}$ mice and controls.**

**(A)** Coefficients for the reinforcer (choice x reward) predictors, fit using trials from predictor blocks separately for *Scn2a*$^{+/-}$ mice and controls.

**(B)** Similar to (A) for choice predictors. The predictor at trial lag of 0 is the bias term.

**(C) – (D)** Similar to (A) – (B) for trials from WSLS blocks. We note the difference between *Scn2a*$^{+/-}$ mice and controls in their reinforcer predictor for a trial lag of 1 (i.e., the last trial) during WSLS blocks. Specifically, control mice had a significant negative coefficient. This means that if the last trial was a rewarded left ($C_{n-1} R_{n-1}$ = -1), then the mouse was more likely to choose right in the current trial because logit $P(C_n) > 0$, whereas if the last trial was a rewarded right ($C_{n-1} R_{n-1}$ = 1), then the mouse was more likely to choose left in the current trial because logit $P(C_n) < 0$. This corroborates with the idea that the control mice were cognizant of the computer opponent's WSLS strategy and countered with a switching behavior. By contrast, this reinforce predictor for a trial lag of 1 was not significantly different from zero for *Scn2a*$^{+/-}$ mice.

Triangle, $P < 0.05$.
